# Gut microbiota promotes pain in fibromyalgia

**DOI:** 10.1101/2023.10.24.563794

**Authors:** Weihua Cai, May Haddad, Rana Haddad, Inbar Kesten, Tseela Hoffman, Reut Laan, Calvin Wong, Nicole Brown, Shannon Tansley, Kevin C. Lister, Mehdi Hooshmandi, Feng Wang, Behrang Sharif, Susan Westfall, Tali Sahar, Charlotte Clayton, Neil Wu, Ji Zhang, Haggai Bar-Yoseph, Milena Pitashny, Jeffrey S. Mogil, Masha Prager-Khoutorsky, Philippe Séguéla, Irah L. King, Yves De Koninck, Nicholas J.B Brereton, Emmanuel Gonzalez, Yoram Shir, Amir Minerbi, Arkady Khoutorsky

## Abstract

Fibromyalgia is a chronic syndrome characterized by widespread pain in the absence of evident tissue injury or pathology, making it one of the most mysterious chronic pain conditions. Despite affecting 2–4% of the population, primarily women^1^, the cause and underlying mechanisms of fibromyalgia remain elusive, and effective targeted treatments are currently unavailable. The gut microbiota of women with fibromyalgia differs from healthy controls^2,3^. However, it is unknown whether changes in gut microbiota have a causal role in mediating pain and other symptoms of fibromyalgia. Here, we show that fecal microbiota transplantation (FMT) from individuals with fibromyalgia, but not from healthy controls, into germ-free mice induces persistent pain hypersensitivity. FMT from fibromyalgia patients led to a reduction in intraepidermal nerve fiber density and alterations in the peripheral immune profile, and induced activation of spinal microglia, which contributed to the development of pain in mice. Notably, the pain hypersensitivity in mice that were administered microbiota from fibromyalgia patients resolved after FMT from healthy controls. Consistent with these findings, an open-label pilot study showed that transplanting microbiota from healthy individuals to humans with fibromyalgia alleviated pain and reduced overall symptom severity. Thus, altered gut microbiota has a causal role in fibromyalgia pain, highlighting it as a promising target for therapeutic interventions.

## Gut microbiota transplantation from fibromyalgia patients induces pain in mice

To investigate whether the gut microbiota in fibromyalgia (FM) patients can cause pain hypersensitivity, we collected fecal samples from women with fibromyalgia and healthy controls (HC) and transplanted them into germ-free mice. For the fibromyalgia group, we selected fecal donors who suffer from primary fibromyalgia, who screened negative for anxiety, depression, irritable bowel syndrome, or other comorbidities (inclusion and exclusion criteria, and demographics are provided in Extended Data Fig. 1a,b), and whose gut microbiota compositional profile was representative of a previously established fibromyalgia signature^2^ (Extended Data Fig. 1c). Fibromyalgia is more prevalent in females, and gut microbiota composition in fibromyalgia has only been studied in women^2–4^; therefore, the weekly FMTs were performed by transplanting human stool from women with fibromyalgia and age-matched healthy controls into germ-free female mice (Fig. 1a). The 16S rRNA gene analysis of feces showed that after the transplantation, diversity differences in the gut microbiota between the two groups of donors were successfully reproduced in recipient mice (Fig. 1b,c, and Extended Data Fig. 2a-c). As expected, significant differences were found between the fecal bacterial composition of fibromyalgia and healthy control microbiota-recipient mice, which increased with time (Fig. 1c; Extended Data Fig. 2c,d; Supplementary Table 1 shows differentially abundant species at week 4 post-FMT). Beta-diversity analysis of whole genome sequencing (WGS) metagenomics data showed substantial differences in the bacterial communities between the two groups, similar to those observed using 16S rRNA gene sequencing (Fig. 1d; total number of sequencing reads, percent of mapping, and microbial composition are provided in Extended Data Fig. 2e-g).

**Fig. 1.**
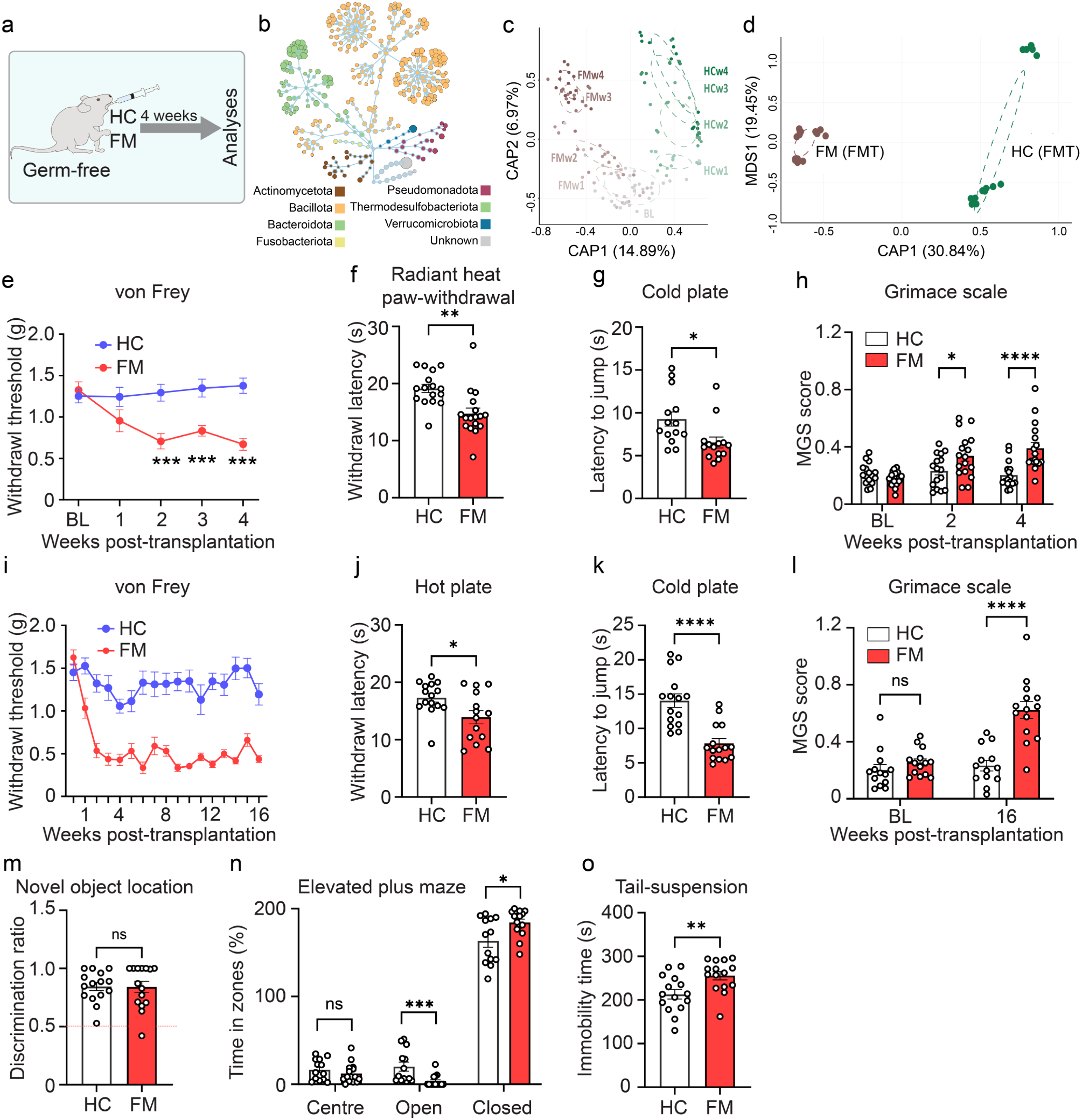
Fecal microbiota transplantation from fibromyalgia patients induces pain hypersensitivity in mice. **a,** Experimental design. Germ-free (GF) female mice received weekly fecal microbiota transplantation (FMT) from healthy controls (HC) and individuals with fibromyalgia (FM). **b**, Flower diagram representing 16S rRNA microbial diversity (1241 ESVs colored by phyla) of mice 4 weeks after transplantation from HC and FM. The size of the distal node (i.e. lowest taxonomic level for an ESV) is proportional to the total raw abundance. **c**, Canonical Analysis of Principal coordinates (CAP; based on Bray-Curtis distances) of normalized (rlog) mouse samples’ ESV abundance at various points (week 1 to 4) after transplantation from HC and FM. Color saturation indicates time whereby darker shades represent later time points. BL: baseline, FMw1-4: week 1-4 post-fibromyalgia FMT, HCw1-4: week 1-4 post-healthy control FMT. **d**, CAP based on Bray-Curtis distances (MDS = multidimensional scaling) of normalized (log) WGS metagenomics contig abundance of mouse samples at week 4 post-transplantation from HC and FM. **e,** Mechanical thresholds were tested weekly in von Frey (HC: n = 30 mice, FM: n = 31 mice), (**f**) heat sensitivity was measured at week 4 post-FMT in radiant heat paw-withdrawal test, (**g**) cold sensitivity was measured at week 4 post-FMT using cold plate test, and (**h**) spontaneous pain was assessed using the Mouse Grimace Scale (MGS) at week 2 and 4 post-FMT (for **f**-**h**; n = 14-17 mice/ group). **i**-**m**, GF mice received HC and FM FMT and were tested for 16 weeks in the (**i**) von Frey test, (**j**) hot plate, (**k**) cold plate, (**l**) MGS, (**m**) novel object location, (**n**) elevated plus maze, and (**o**) tail-suspension tests (for **i**-**o;** n = 13-15 mice/ group). Two-tailed, unpaired Student *t*-tests in **f**,**g**,**j**,**k**,**m**,**o**. Two-way ANOVA followed by Tukey’s post-hoc comparison in **e**,**h**,**i**,**l**,**n**. \**P* < 0.05, \*\**P* < 0.01, \*\*\**P* < 0.001, \*\*\*\**P* < 0.0001; ns – not significant. All data are presented as mean ± s.e.m.

Behavioral studies showed that FMT from individuals with fibromyalgia, but not healthy controls, into germ-free mice induced the development of mechanical hypersensitivity (in the von Frey test, Fig. 1e), thermal hypersensitivity (in the radiant heat paw-withdrawal test, Fig. 1f), cold hypersensitivity (in the cold-plate test, Fig. 1g) and spontaneous pain (measured via the Mouse Grimace Scale [MGS], Fig. 1h) at week 4 post-transplantation. Chronic widespread pain in fibromyalgia patients is often accompanied by cognitive symptoms and depression^5,6^. Mice transplanted with fibromyalgia gut microbiota did not show deficits in cognitive functions 4 weeks post-FMT, compared to mice transplanted with gut microbiota from healthy individuals. No change was found in memory (in the novel object location [NOL] test, Extended Data Fig. 3a), anxiety (in the elevated plus maze [EPM], Extended Data Fig. 3b), or depression-like behavior (in the tail-suspension test, Extended Data Fig. 3c). We found no evidence of intestinal barrier dysfunction or gut inflammation in fibromyalgia microbiota-recipient mice 4 weeks after fecal transplantation as indicated by unchanged histomorphology (Extended Data Fig. 4a), gut permeability (using FITC-dextran 4 kDa, Extended Data Fig. 4b), tight-junction gene levels (Occludin [*Ocln*], Zonula Occludens-1 and -2 [*Zo1*/*Zo2*], Extended Data Fig. 4c) and body weight (Extended Data Fig. 4d).

An additional cohort of germ-free mice was colonized with fibromyalgia and healthy control microbiota and monitored for 4 months to study the long-term effects. Four months after the transplantation, the mice that received fibromyalgia FMT showed mechanical (Fig. 1i), heat (Fig. 1j) and cold (Fig. lk) hypersensitivity, and spontaneous pain (Fig. 1l). No memory deficits (Fig. 1m) were observed. However, unlike at the 4-week time-point, fibromyalgia microbiota-recipient mice spent less time in the open arms in the EPM (Fig. 1n) and exhibited increased immobility in the tail-suspension test (Fig. 1o), suggesting the development of a depression-like phenotype. Taken together, these results show that gut microbiota from women with fibromyalgia is sufficient to induce persistent pain in mice and, over an extended period, can lead to the development of a depression-like phenotype.

We also performed FMT from fibromyalgia patients and healthy controls into specific-pathogen-free (SPF) mice that were pre-treated with antibiotics for one week prior to transplantation (Extended Data Fig. 5a). While we observed mechanical and heat hypersensitivity, but not spontaneous pain, following microbiota transplantation from fibromyalgia patients (Extended Data Fig. 5b-g), the hypersensitivity returned to baseline levels 5 weeks post-FMT. This suggests that this approach may not be optimal for studies of chronic pain in fibromyalgia.

## Pervasive changes in mice colonized with fibromyalgia microbiota

Gut microbiota can affect metabolomic profile and immune functions, both of which are dysregulated in fibromyalgia patients^7–10^. Using untargeted metabolomics analysis, we found that mice that received fibromyalgia FMT exhibited an altered metabolomic profile as compared to mice that received FMT from healthy controls. We detected changes in the metabolism of neuroactive amino acids, including increased levels of glutamine in the spinal cord and elevated glutamate in the brain (Extended Data Fig. 6; Supplementary Table 2). Lipid metabolism was also dysregulated in fibromyalgia FMT-recipient mice, as they exhibited a decrease in medium and long**-**chain fatty acids, branched-chain fatty acids and dicarboxylate fatty acids in plasma (Extended Data Fig. 6). Moreover, we found alterations in secondary bile acid metabolism (decreased ursocholate in plasma and increased lithocholic acid sulphate in the feces), consistent with previous findings in fibromyalgia^4^.

An analysis of the immune landscape, conducted using single-cell RNA sequencing (scRNA-seq) of peripheral blood mononuclear cells (PBMCs) (Fig. 2a and Extended Data Fig. 7a,b), revealed that mice colonized with fibromyalgia microbiota demonstrate an increased proportion of classical monocytes compared to mice that received healthy control FMT (Fig. 2b,c). Analysis of differentially expressed genes within this cellular population showed enrichment for IL-17 and TNF signaling pathways, which are associated with inflammatory responses (Fig. 2d). Immune-related changes were also present in other cell types, including intermediate/non-classical monocytes, memory/plasma B cells, DC, Treg, and B cell-like T cells (Supplementary Table 3). The proportion of memory B cells was decreased in fibromyalgia FMT-recipient mice (Extended Data Fig. 7c).

**Fig. 2.**
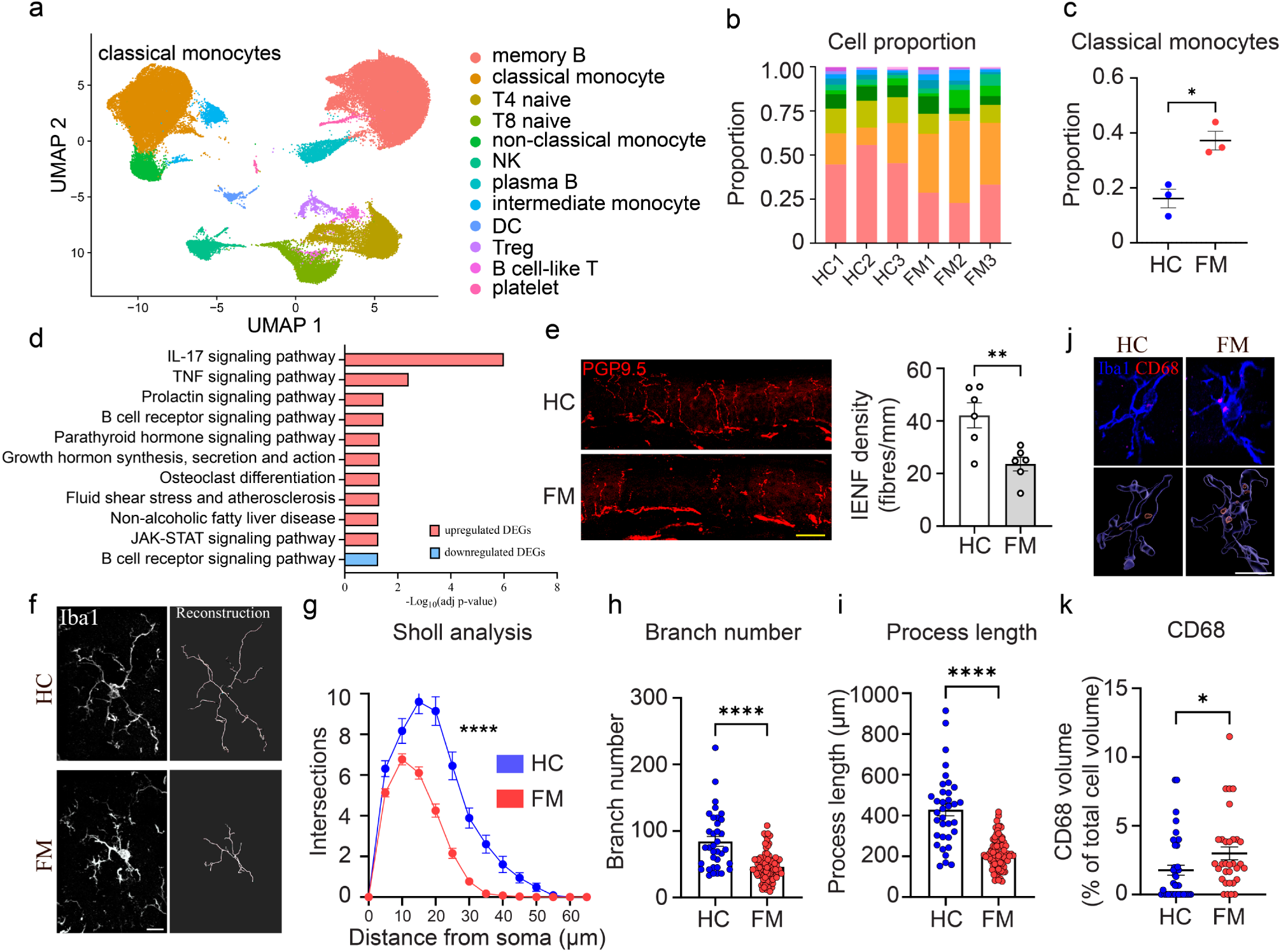
Fecal microbiota transplantation from fibromyalgia patients induces changes in various systems in mice. Peripheral blood mononuclear cells (PBMCs) of mice that received microbiota from HC and FM individuals were collected at week 4 post-transplantation (n = 3 mice/group). **a,** UMAP plot showed 12 distinct subpopulations, different colors correspond to different cell types. **b,** The columns show the proportion of different cell types in each sample. **c**, The proportion of classical monocytes was increased in the FM group. **d**, Analysis of differentially expressed genes (DEGs) in classical monocytes revealed changes in distinct pathways (upregulated genes (red) and downregulated genes (blue)). The y-axis indicates the representative KEGG pathway terms. **e**, Immunohistochemistry of glabrous hind paw skin (week 4 post-FMT) using anti-PGP9.5 antibody revealed decreased density of intraepidermal nerve fibers (IENF) (n = 6 mice/group). **f,** Iba1^+^ microglia (left) and 3D reconstructions (right) in HC-and FM-recipient mice (week 4 post-FMT) lumbar dorsal spinal cord. **g,** Sholl analysis of reconstructions (HC: n = 7 mice/35 cells, FM: n = 7 mice/78 cells for **g**-**i**). (**h**,**i**) Reduced microglia branch number and total process length. **j,** Immunofluorescence images (upper) and 3D volume renders (bottom) of CD68 staining in microglia. **k**, Quantification of CD68 as a percentage of the total cell volume (HC: n = 7 mice/41 cells, FM: n = 7 mice/32 cells). Scale bars; **e**, 20 μm; **f,j**. 10 μm. Two-tailed, unpaired Student *t*-tests in **c**,**e**,**h**,**i**,**k**. Two-way ANOVA followed by Tukey’s post-hoc comparison in **g**. \**P* < 0.05, \*\**P* < 0.01, \*\*\*\**P* < 0.0001. All data are presented as mean ± s.e.m.

A subset of individuals with fibromyalgia exhibits reduced intraepidermal nerve fiber density^11–13^. Imaging of PGP9.5-positive nerve fibers in the glabrous skin of mice that received fibromyalgia FMT revealed reduced density of sensory fibers in the epidermis as compared to healthy control FMT-recipient mice (Fig. 2e). Collectively, these results show that the transplantation of fibromyalgia microbiota into germ-free mice induces multisystemic effects, including altered metabolic profile, low-grade inflammation, and reduced epidermal innervation.

## Activation of microglia contributes to hypersensitivity

Low-grade peripheral inflammation observed in mice that received fibromyalgia gut microbiota prompted us to evaluate microglia, the principal immune cells in the central nervous system, which are often activated in pathological pain states and contribute to the development of pain hypersensitivity^14^. In fibromyalgia FMT-recipient mice, microglia were present in a reactive state in the lumbar dorsal spinal cord as evident by morphological changes (decreased number of intersections across the arbor in 3D Sholl analysis, and reduced branch number and length) (Fig. 2f-i), and the increased expression of lysosomal CD68, a marker of microglial phagocytic activity (Fig. 2j). Reactive microglia were also found in other areas of the spinal cord (lumbar ventral, and thoracic dorsal and ventral horns, Extended Data Fig. 8a-l) and in the anterior cingulate cortex (ACC) (Extended Data Fig. 8m-p). To study the functional role of microglia in mediating pain phenotypes, we depleted microglia by administering an inhibitor of colony-stimulating factor 1 receptor (CSF1R), PLX5622, for 7 days prior to transplantation (Fig. 3a). Colonization of microglia-depleted germ-free mice with fibromyalgia microbiota induced mechanical and cold hypersensitivity that was lower than in mice with intact microglia (Fig. 3b,c). Heat hypersensitivity and spontaneous pain were not affected by microglia depletion (Extended Data Fig. 9a,b). These results indicate that the development of pain after fibromyalgia FMT is partially mediated by microglia.

**Fig. 3.**
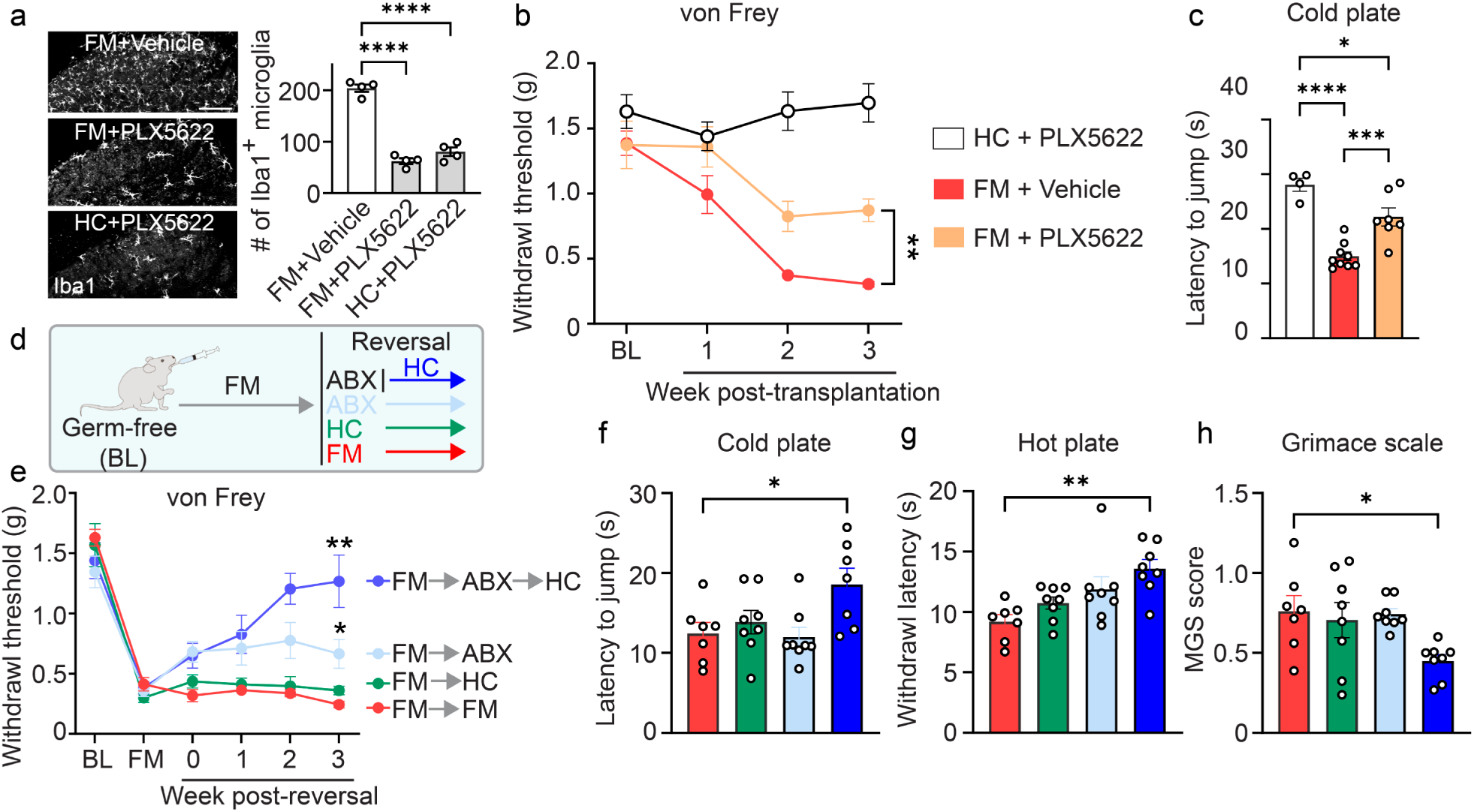
Alleviation of pain hypersensitivity via targeting microglia or substituting microbiota. Germ-free (GF) mice were treated with PLX5622 before receiving fecal microbiota transplantation (FMT) from HC and FM patients. **a**, Immunostaining for Iba1 shows a reduction in the number of microglia in the dorsal horn following PLX5622 treatment, n = 4 mice/group. Scale bar, 100 μm. Transplantation of microbiota from fibromyalgia patients (FM FMT) in microglia-depleted mice induced (**b**) less mechanical hypersensitivity and (**c**) less cold hypersensitivity as compared to vehicle-treated animals (HC+PLX5622: n = 4 mice; FM+Vehicle: n = 9 mice; FM+PLX5622: n = 7 mice). **d**, Experimental design of reversal experiment using transplantation of microbiota from healthy control donors (HC FMT). Mice were randomly divided into 4 groups after four weeks of FM FMT: FM-ABX-HC (blue, n = 8 mice) received one week of ABX pretreatment (and PEG on the last day) followed by HC FMT weekly; FM-ABX (light blue, n = 8 mice) were continuously treated with ABX; FM-HC (brown, n = 8 mice) received HC FMT weekly without ABX/PEG pretreatment; FM-FM (red, n = 7 mice) continued to receive weekly FM FMT. **e,** Mechanical thresholds were tested weekly in all groups and (**f**) cold sensitivity, (**g**) heat sensitivity, and (**h**) spontaneous pain were tested at week 3 post-reversal. BL: baseline, time before transplantation; FM: 4 weeks after fibromyalgia microbiota transplantation; ABX: antibiotics treatment. One-way ANOVA followed by Tukey’s post-hoc comparison in **a**,**c**,**f**,**g**,**h**. Two-way ANOVA followed by Tukey’s post-hoc comparison in **b**,**e**. \**P* < 0.05, \*\**P* < 0.01, \*\*\**P* < 0.001, \*\*\*\**P* < 0.0001. All data are presented as mean ± s.e.m.

## The effect of fibromyalgia microbiota on pain can be reversed by microbiota from healthy individuals

We next tested a therapeutically oriented approach by substituting fibromyalgia microbiota with the microbiota from healthy donors. Germ-free mice were first administered fibromyalgia FMT, and four weeks later, when the pain hypersensitivity had fully developed, were treated with an antibiotic cocktail for one week (and polyethylene glycol [PEG] on the last day) to deplete their microbial communities, followed by FMT from healthy controls (experimental design shown in Fig. 3d). Transplantation of healthy control microbiota into fibromyalgia microbiota-recipient mice, after antibiotic pre-treatment, induced a shift in the bacterial composition of the gut microbiota (Extended Data Fig. 10a,b) and this shift was diminished if antibiotics were omitted. Behavioral analysis showed that re-colonization of fibromyalgia microbiota-recipient mice with FMT from healthy donors reversed the mechanical hypersensitivity (Fig. 3e) and alleviated cold (Fig. 3f) and heat (Fig. 3g) hypersensitivity, and spontaneous pain (Fig. 3h). Notably, healthy control FMT into fibromyalgia microbiota-recipient mice without prior suppression of gut communities with antibiotics failed to alleviate pain (Fig. 3e-h). Antibiotic treatment alone slightly reduced mechanical hypersensitivity but had no effect on heat and cold hypersensitivity or spontaneous pain (Fig. 3e). No differences in body weight between the groups were found (Extended Data Fig. 10c). These results show that substitution of fibromyalgia-associated bacterial communities with microbiota from healthy donors reverses pain hypersensitivity.

## FMT from healthy individuals improves fibromyalgia symptoms in humans

Building on our findings in mice, we conducted an open-label, pilot study to investigate the effect of FMT from healthy individuals on the symptoms of fibromyalgia in humans. Fourteen women (age 51±13.7 years; demographics are provided in Extended Data Fig. 11a) with severe, refractory fibromyalgia were enrolled. Despite receiving the usual clinical care, all participants reported significant symptomatic burden with high pain intensity and profuse fatigue (fibromyalgia-related variables are shown in Extended Data Fig. 11b). Quantitative sensory testing (QST) showed pain hypersensitivity in response to heat and cold stimuli in 54% and 23% of patients, respectively (QST data are provided in Extended Data Fig. 12). Following depletion of the endogenous microbial communities using antibiotics and bowel cleansing, each patient received five FMTs, once every two weeks, via oral administration of encapsulated transplants from healthy donor (age 34.7 ± 14.2 years; treatment regimen is described in Methods and shown in Fig. 4a; laboratory tests for donor screening are shown in Extended Data Fig. 13a). The treatment was overall well tolerated; adverse effects included mainly increased fatigue and mild gastrointestinal symptoms (Extended Data Fig. 13b). One week after the last FMT, pain intensity decreased by ≥2 points (out of 10, using pain numerical rating scale) in 12 of the 14 participants, resulting in a clinically and statistically significant decrease in the mean pain intensity (Fig. 4b,c). Overall, the symptomatic burden decreased, and anxiety, depression, sleep quality, and physical quality of life scores improved (Fig. 4d-h), while mental quality of life scores showed a trend toward improvement (p=0.07, Fig. 4i). Following FMT from healthy donors, QST measures showed reduced cold pain hypersensitivity (Fig. 4j and Extended Data Fig. 12) and a trend toward improved heat pain hypersensitivity (p=0.07, Fig. 4k), whereas measures of conditioned pain modulation (CPM) and detection thresholds of non-noxious thermal stimuli (warm and cold) were unchanged (Extended Data Fig. 12).

**Fig. 4.**
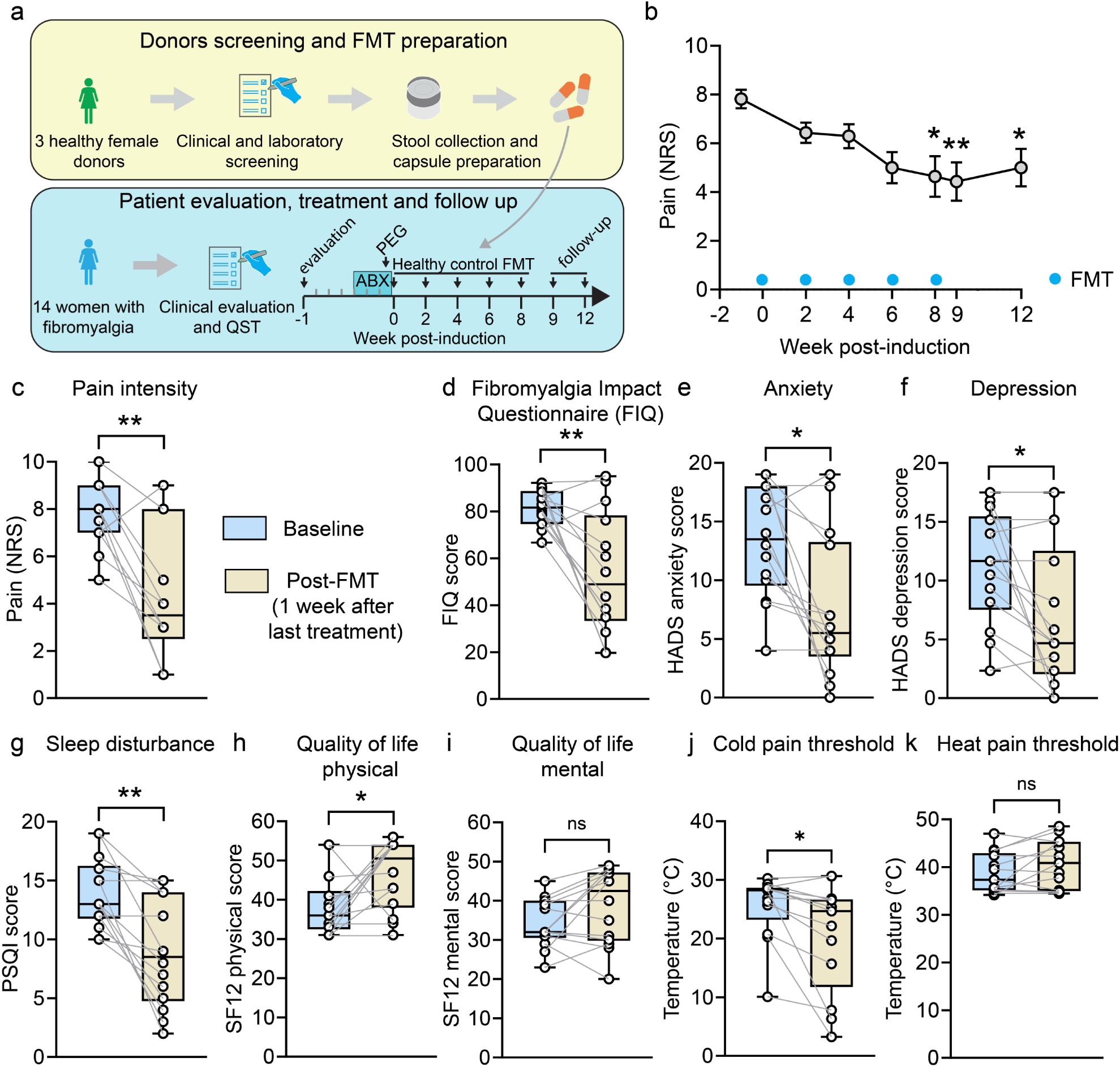
Open-label clinical trial of healthy control FMT in humans with fibromyalgia. **a,** Schematic representation of the study design. **b**, Pain intensity over time (Numeric Rating Scale [NRS], presented as mean ± s.e.m, Friedman One-Way Repeated Measure Analysis of Variance, *adjusted *P* < 0.05, **adjusted *P* < 0.01). FMT treatments are marked by blue dots. **c,** Pain intensity at baseline and up to four weeks post-treatment (NRS). **d**, Overall symptom severity as measured by the Fibromyalgia Impact Questionnaire (FIQ) at baseline and one week post-treatment. **e-f**, HADS (hospital anxiety and depression scale) anxiety and depression scores at baseline and one week post-treatment. **g,** Sleep disturbance scores as measured by the PSQI (Pittsburgh Sleep Quality Index) at baseline and one week post-treatment. **h-i**, Physical-and mental quality of life as measured by the SF-12 questionnaire at baseline and one-week post-treatment. **j-k**, QST measures of cold and heat pain thresholds at baseline and one week post-treatment (n = 14 women). In **c**-**k,** Wilcoxon signed-rank test was used, \*\**P*<0.01, **\****P*<0.05. Data in **c**-**k** are presented as a box and whisker plot.

## Discussion

We found that the transplantation of gut microbiota from fibromyalgia patients induces pain in mice. This finding suggests that the altered gut microbiota in fibromyalgia patients may play a causal role in the disease’s pathophysiology, contributing to widespread pain. This conclusion is bolstered by clinical findings from an open-label pilot study, in which women with fibromyalgia experienced clinically significant improvements in their pain levels and overall symptoms after FMT from healthy individuals.

The colonization of germ-free mice with the gut microbiota from individuals with fibromyalgia impacted numerous systems as compared to FMT from healthy controls. We observed alterations in the metabolomic profile, particularly in glutamate, lipid, and bile acid metabolism, and detected an increase in peripheral monocytes and the activation of spinal microglia. Notably, a CSF1R inhibitor, which targets circulating and tissue macrophages^15^, including microglia, only partially alleviated the pain hypersensitivity, suggesting the additional involvement of other mechanisms. Furthermore, we found a decrease in the intraepidermal innervation of fibromyalgia microbiota-recipient mice. The functional role of reduced density of intraepidermal fibers, which has been linked to immunological and metabolic disturbances, in mediating pain and other phenotypes in fibromyalgia is not well understood^16–19^. Together, our findings highlight the complex and multifactorial effects of fibromyalgia microbiota across several systems in mice. Similarly, humans with fibromyalgia exhibit a wide range of phenotypes, including the sensitization of the peripheral and central nervous systems^20,21^, immune dysregulation^22–24^, and metabolomic alterations^3,25–29^. Notably, individuals with fibromyalgia often experience functional gastrointestinal disorders, including irritable bowel syndrome (IBS) and functional dyspepsia^30,31^. Our results raise the intriguing possibility that dysregulation in gut microbiota-host interactions in fibromyalgia patients contributes to various molecular and behavioral phenotypes, likely through diverse mechanisms that require further elucidation.

Multiple environmental factors have been suggested as potential triggers for fibromyalgia, including psychosocial stressors, infections, and inflammatory diseases^32^. Additionally, environmental and host-related factors, such as diet, stress and physical activity, are also known to affect the severity of symptoms in humans with fibromyalgia. Interestingly, these and other environmental factors can induce long-lasting alterations in the composition of the gut microbiota in humans^33^. It is, therefore, compelling to speculate that the gut microbiota could mediate some of the environmental effects in fibromyalgia.

Our study in humans demonstrated a positive clinical effect of FMT from healthy individuals to fibromyalgia patients. In addition to patient self-report of reduced pain, we also measured improved cold pain thresholds, providing a more objective assessment of the treatment’s effect. While our clinical study is limited by patients not being blinded to the treatment and by the lack of a control arm, it represents the first evidence of beneficial therapeutic effects of FMT in fibromyalgia. We believe that these promising results warrant further examination of this strategy.

FMT from human donors with fibromyalgia into mice induced depression-like behavior at 4 months but not 4 weeks after transplantation. These results may suggest that, in this study, the depression phenotype is not directly transferred through the gut microbiota but likely emerges as a consequence of prolonged pain hypersensitivity. This conclusion is consistent with the selection of human donors without anxiety and depression as well as with previous studies showing that chronic pain can lead to the development of anxiety and depression-like behavior within several months^34,35^.

Recent advances in understanding the key functions of the gut microbiota and the gut-brain axis in various disorders of the nervous system have facilitated the development of microbial-based interventions, such as probiotics, FMT, or treatment with bacterial-derived metabolites. Our study, which establishes the functional significance of the gut microbiota in fibromyalgia, should facilitate the evaluation of such therapeutic approaches for this prevalent chronic pain syndrome.

**Extended Data Fig. 1.**
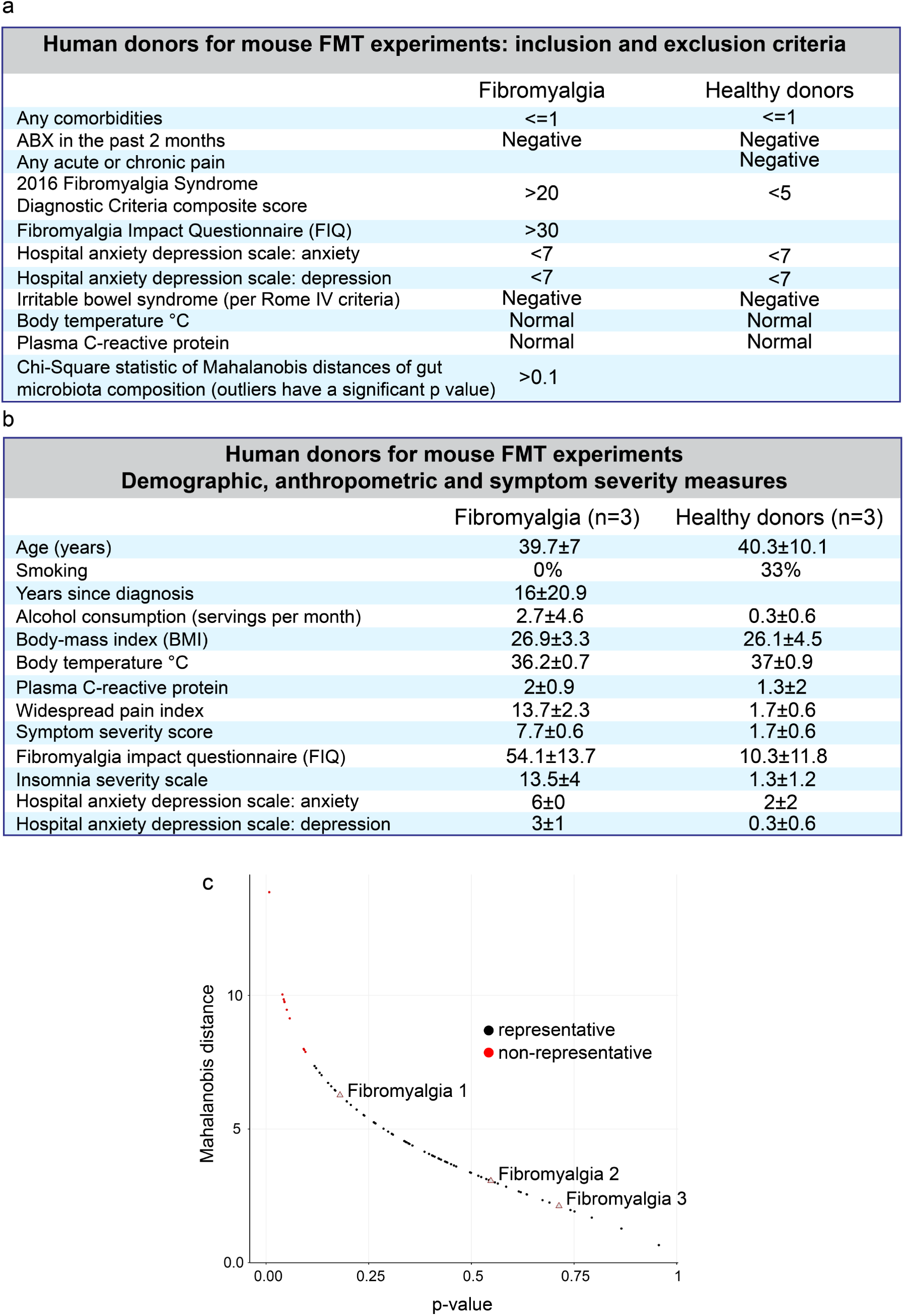
Inclusion criteria, exclusion criteria, and demographics for donor selection. **a,** Inclusion and exclusion criteria of human donors for mouse FMT experiments. **b**, Demographic, anthropometric, and symptom severity measures of human donors for mouse FMT experiments. **c**, Mahalanobis distances of 78 samples (16S rRNA ESV table of FM patient stool microbiome) as a function of significance of the Chi-Square statistic of Mahalanobis distances (represented as a p-value). Samples with a p-value >0.1 were not significant (i.e., they represented outliers) and represented in red. The 3 selected samples are represented by triangles.

**Extended Data Fig. 2.**
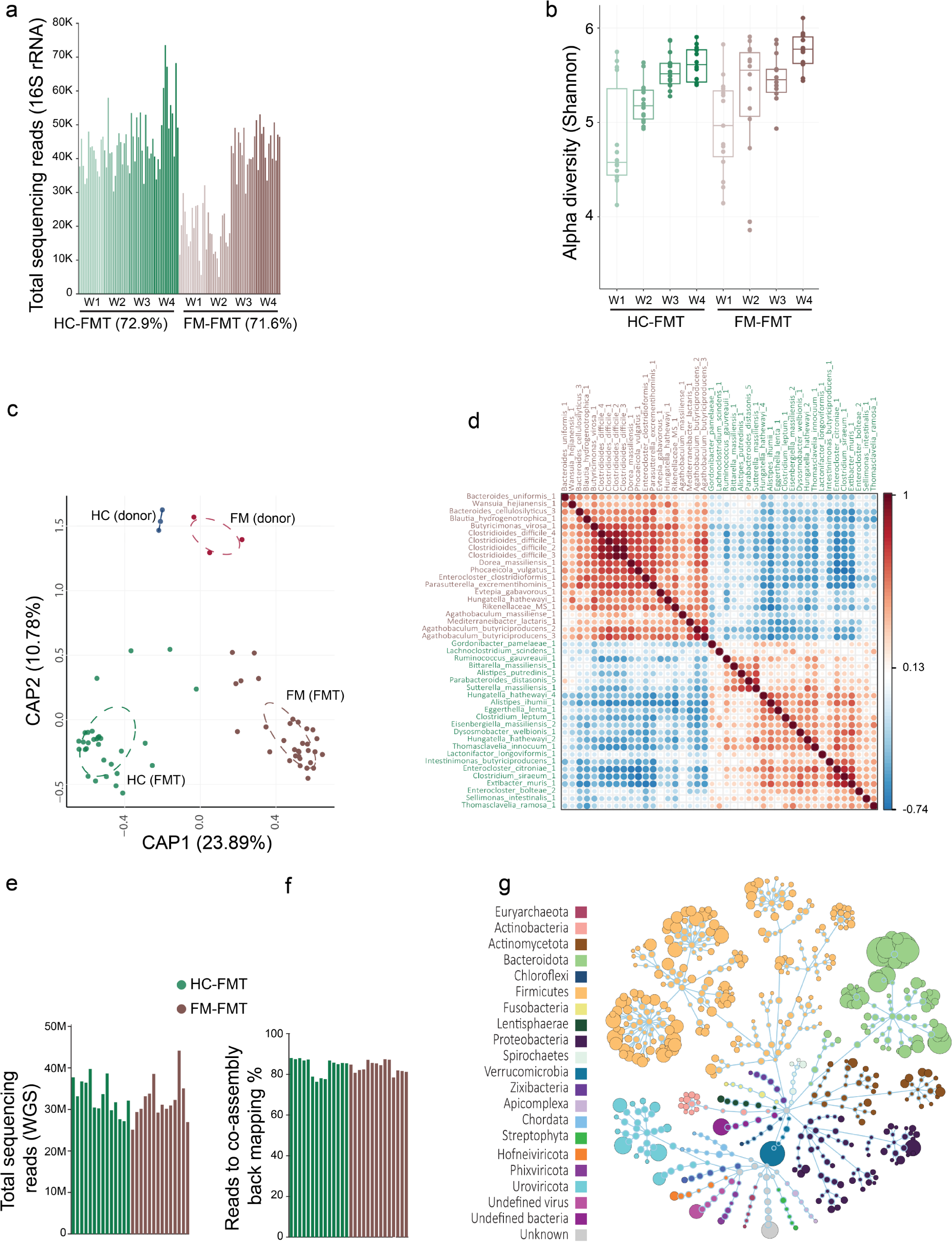
Characterization of gut microbiota. **a,** 16S rRNA library sizes from different samples (MiSeq sequencing). The number in brackets represents the percentage of raw reads that was used to infer ESV during ANCHOR pipeline. **b,** Comparison of alpha diversity indices (Shannon). **c,** CAP based on Bray-Curtis distances of normalized (rlog) mouse (week 4 after transplantation; labeled HC/FM (FMT)) and human donor samples’ ESV abundance (labeled HC/FM (donor)). **d,** ESV correlation (normalized abundance; rlog) using Kendall rank heatmap. Colors in label represent differentially abundant ESVs (purple: higher in FM group; green: higher in HC group). DESeq2 was used for differential abundance analysis with a significance threshold of 0.1 FDR. **e,** WGS library sizes from fecal samples of recipient mice (week 4 post-FMT). **f,** Back-mapping rate for each sample using initial raw reads and the co-assembly. **g,** Microbial diversity (represented as a flower diagram) of all co-assembly contigs with annotation (alignment minimum identity: 90%). The size of the distal node (i.e. lowest taxonomic level for a contig) is proportional to the total raw abundance.

**Extended Data Fig. 3.**
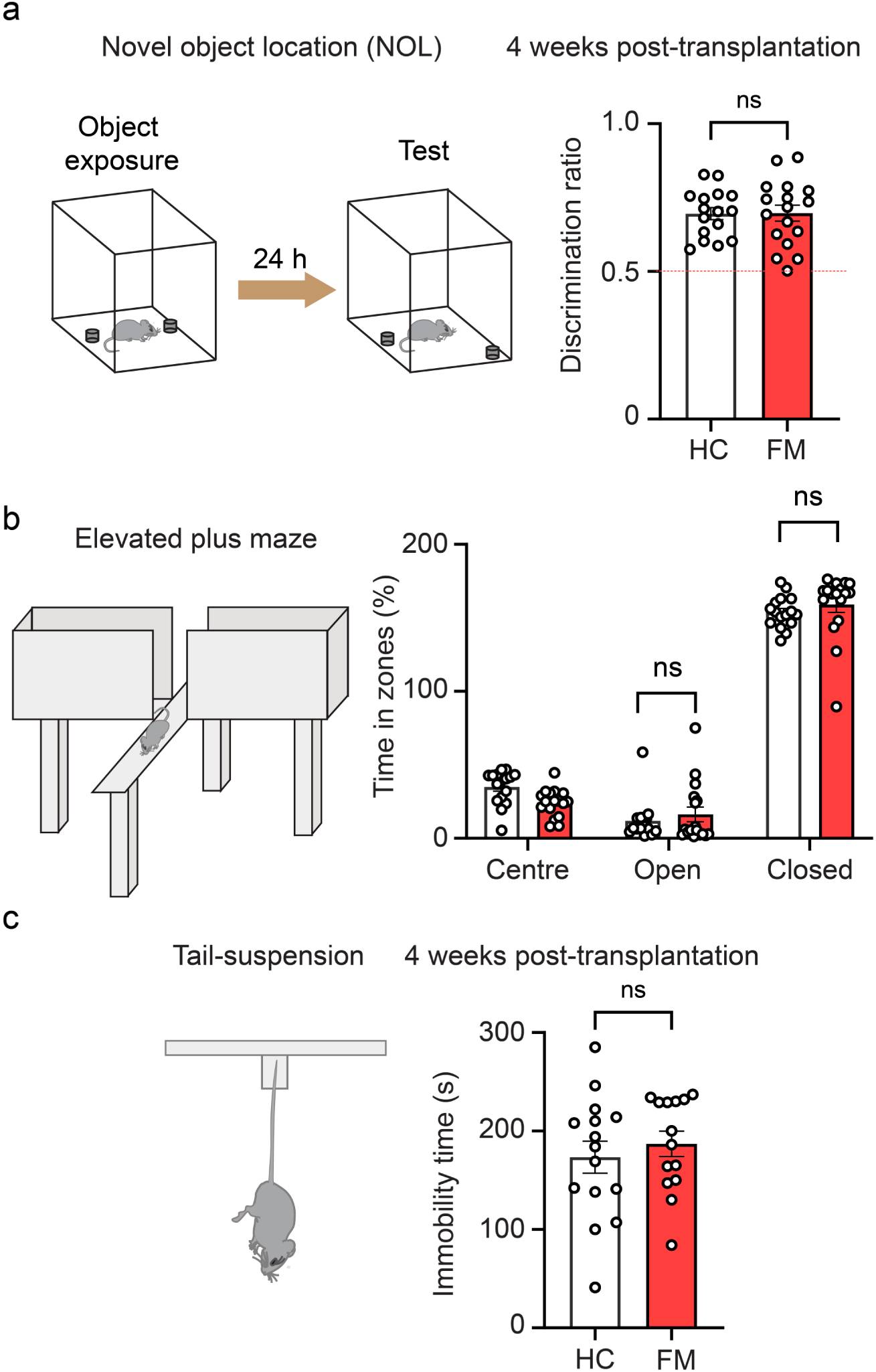
FMT from fibromyalgia patients does not affect cognitive functions in mice at week four post-transplantation. Germ-free mice received weekly FMT from HC and individuals with FM. Four weeks after the first FMT, the mice were tested in the (**a**) novel object location (NOL), (**b**) elevated plus maze (EPM), and (**c**) tail-suspension tests (n = 14-17 mice/group). Two-tailed, unpaired Student *t*-tests in **a,c**. One-way ANOVA followed by Tukey’s post-hoc comparison in **b**. ns – not significant. All data are presented as mean ± s.e.m.

**Extended Data Fig. 4.**
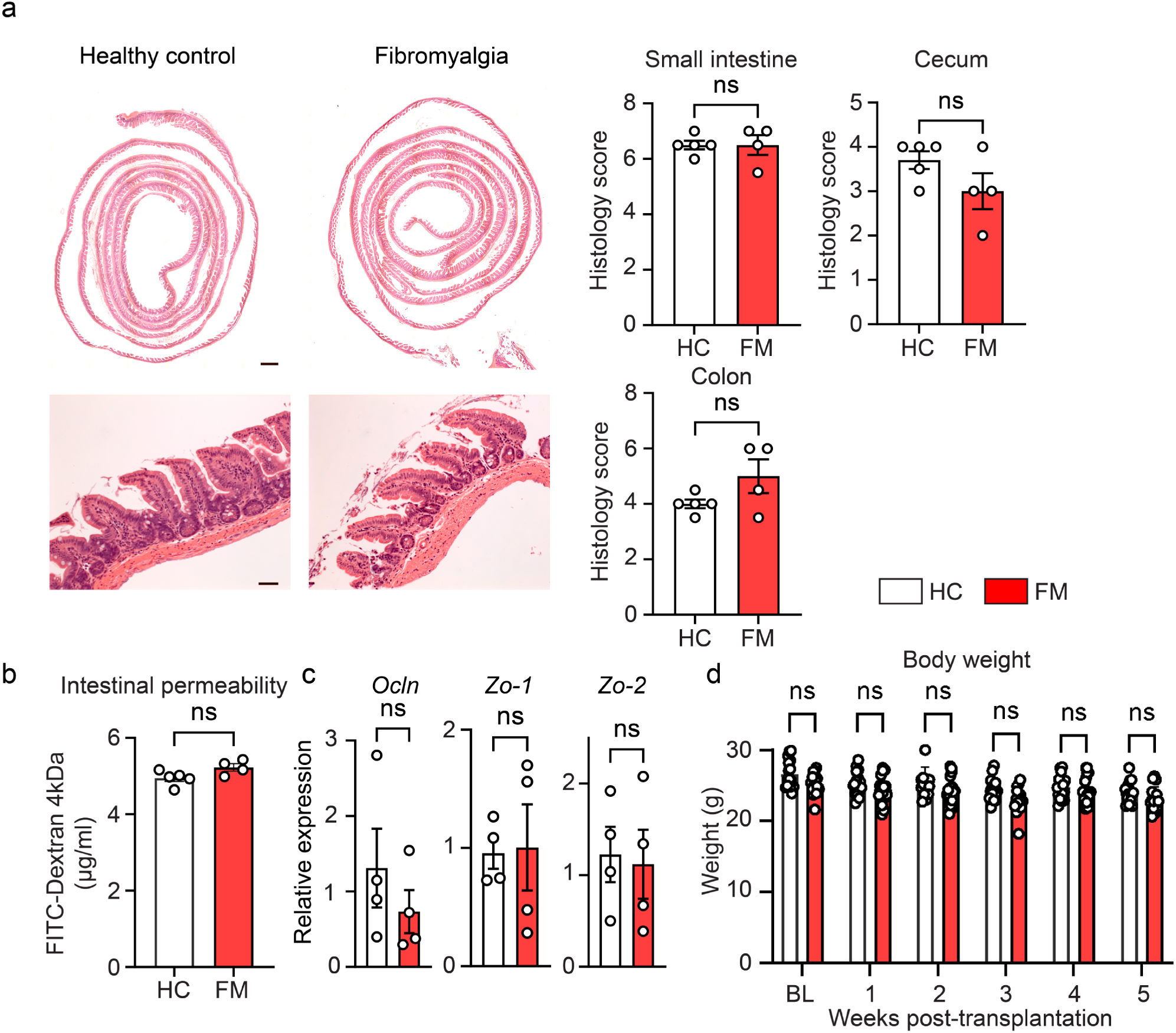
FMT from fibromyalgia patients does not induce gut inflammation in recipient mice. Gastrointestinal characterization of mice at week 4 post-FMT. **a**, H&E staining of small intestine. upper: ’Swiss roll’, scale bar, 500 μm; bottom: scale bar, 50 μm. Histologic evaluation revealed no differences between the groups (n = 4 - 5 mice/group). **b**, Intestinal permeability using FITC Dextran (4 KDa) in serum following oral gavage (n = 4 - 5 mice/group). **c**. Expression of the tight-junction genes Occludin (*Ocln*), Zonula Occludens 1 (*Zo1*), and Zonula Occludens 2 (*Zo2*) in the small intestine of FM and HC FMT-recipient mice, as measured by qRT-PCR (n = 4 mice/group). **d**, Mouse body weight (n = 16 - 17 mice/group). BL: time point before FMT. Two-tailed, unpaired Student *t*- tests in **a,b,c**. Two-way ANOVA followed by Tukey’s post-hoc comparison in **d**. ns – not significant. All data are presented as mean ± s.e.m.

**Extended Data Fig. 5.**
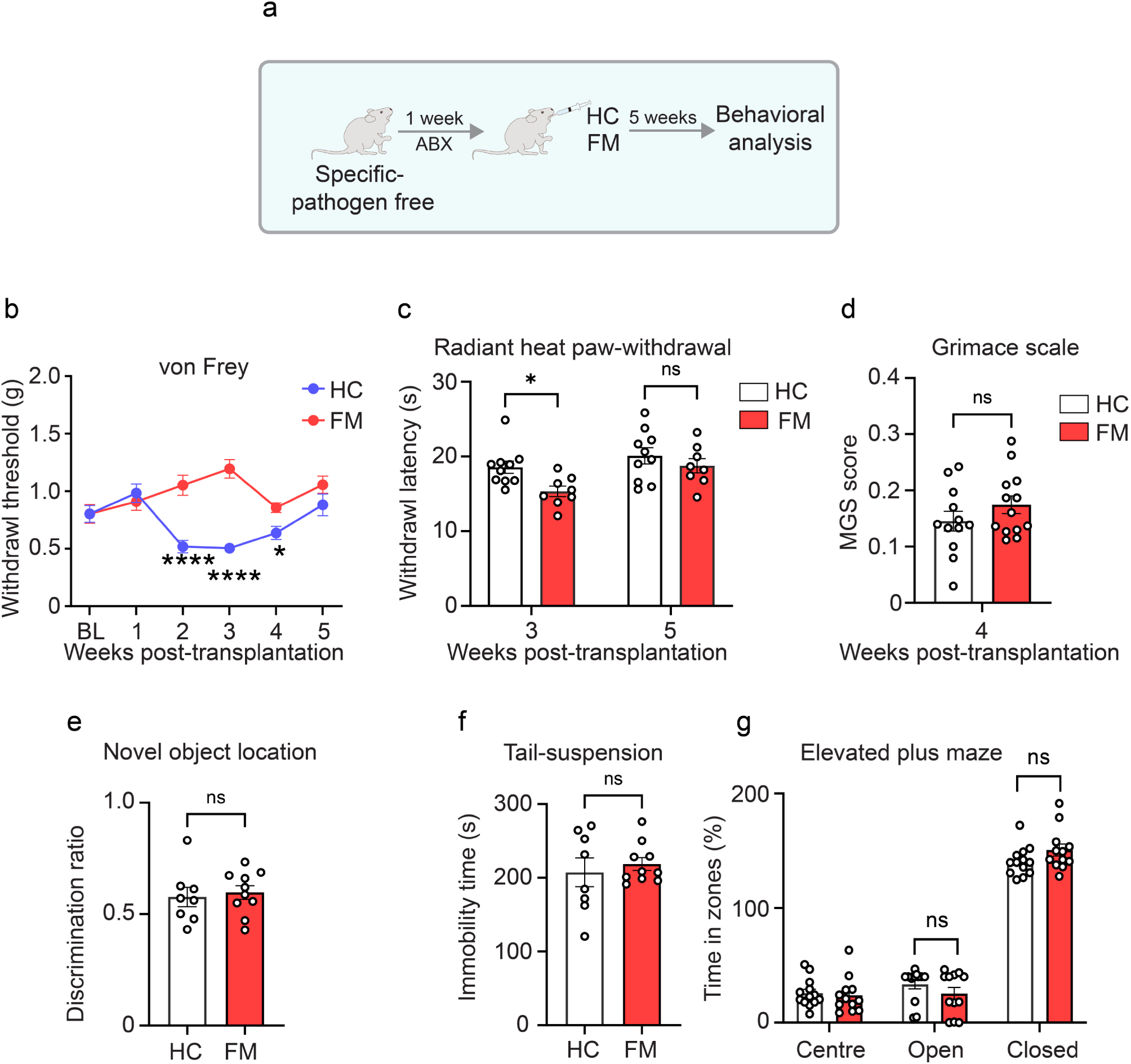
FMT from fibromyalgia patients induces transient pain hypersensitivity in antibiotics-treated specific pathogen-free (SPF) mice. Specific pathogen-free (SPF) female mice were treated with antibiotic cocktail in drinking water for one week (see methods for details) and then received weekly FMT from HC and individuals with FM for 5 weeks. **a**, A schematic illustration of the experimental design. **b,** Mechanical thresholds in von Frey test were assessed weekly, (**c**) heat thresholds in radiant heat paw-withdrawal test were measured at week 3 and 5 post-FMT, and (**d**) spontaneous pain in the Mouse Grimace Scale (MGS) was tested at week 4 post-FMT. At week 4 post-FMT, cognitive functions were assessed, including (**e**) memory in the novel object location test, (**f**) depression-like behavior in the tail-suspension test, and (**g**) anxiety in the elevated plus maze (n = 8 - 13 mice/group). Two-tailed, unpaired Student *t*-tests in **d**,**e**,**f**. Two-way ANOVA followed by Tukey’s post-hoc comparison in **b**,**c**,**g**. \**P* < 0.05; ns – not significant. All data are presented as mean ± s.e.m.

**Extended Data Fig. 6.**
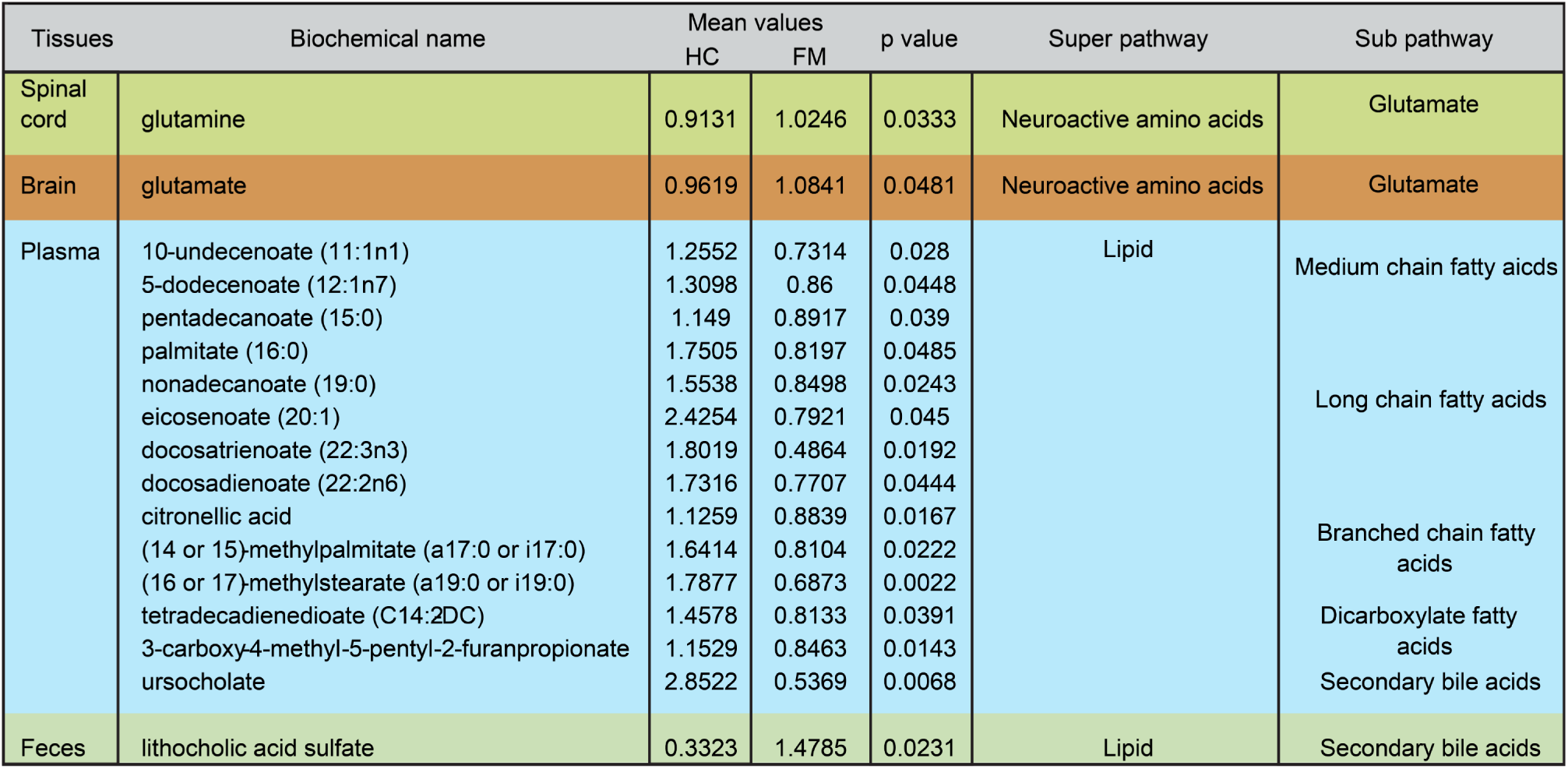
Metabolomics changes after fecal microbiota transplantations in mice. Changes in metabolites in the spinal cord, brain, plasma, and feces samples of mice at week 4 post-FMT from HC and FM patients (complete dataset is provided in Supplementary Table 2). Biochemical name, mean values, p-value, super metabolomic pathways and sub metabolomic pathways are shown. Differences in means were tested by Welch’s two-sample *t*-test. n = 5 mice/group, FM vs HC FMT-recipient mice.

**Extended Data Fig. 7.**
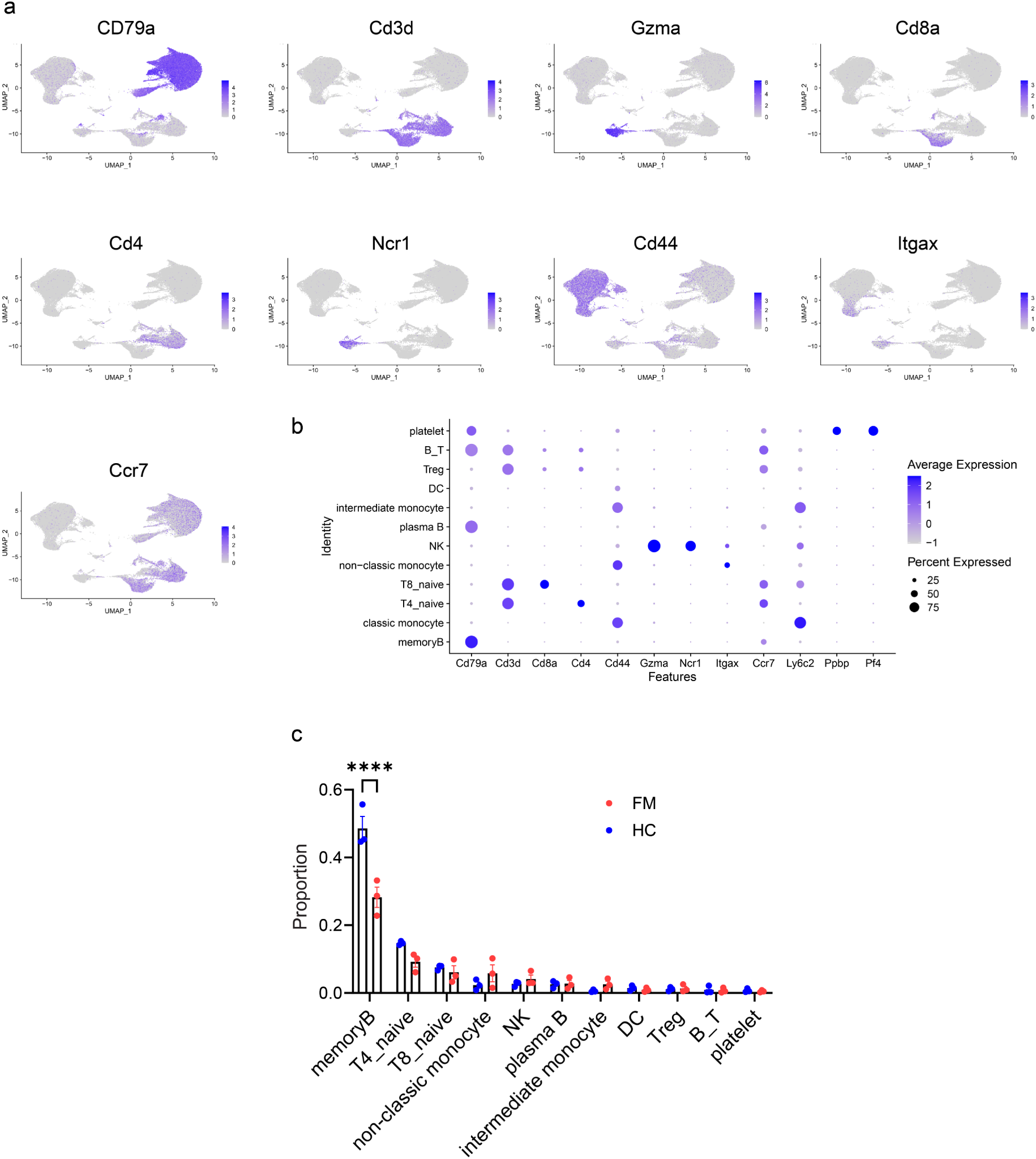
Characterization of immune cells using scRNAseq. Mice received microbiota from HC and FM patients and peripheral blood mononuclear cells (PBMC) were collected at week 4 post-transplantation (n = 3 mice/group), followed by scRNAseq analysis. **a**, Expression of nine representative genes with differential expression, shown using UMAP plots in Fig. 2a. **b**, Dot plot exhibits markers specifically expressed for each cluster. The size of the dots indicates the proportion of the cells that express that gene within the cluster. The color of the dots corresponds to the expression level of that gene. **c**, Columns display the relative percentage of immune cell types in PBMCs from FM and HC groups (except classical monocyte). Two-way ANOVA followed by Tukey’s post-hoc comparison in c. \*\*\*\**P* < 0.0001. All data are presented as mean ± s.e.m.

**Extended Data Fig. 8.**
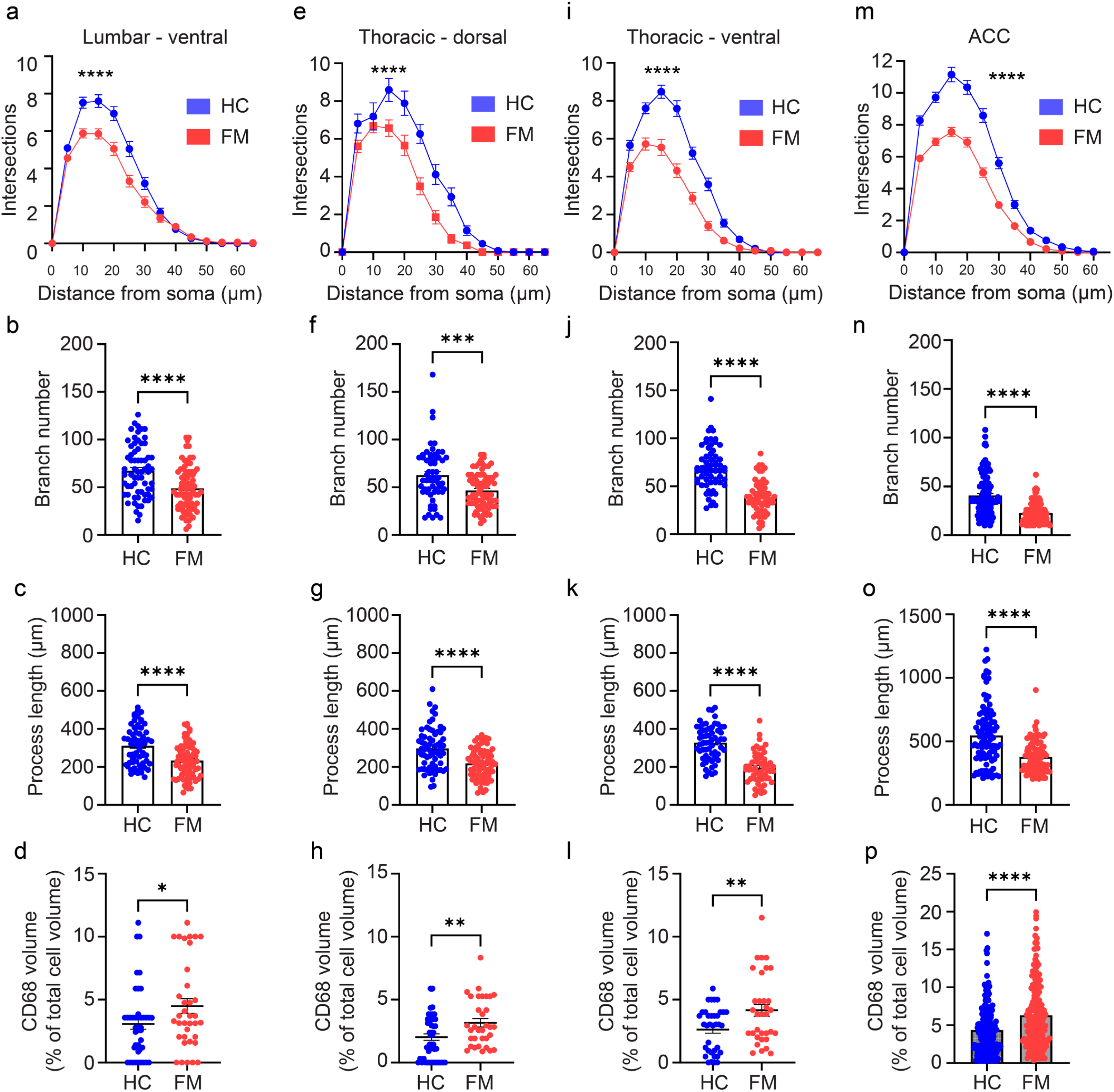
Altered microglia in mice following fibromyalgia FMT. Germ-free mice received weekly FMT from HC and individuals with FM. Four weeks after the first FMT, microglia in the lumbar dorsal horn were immunostained for Iba1 and CD68. Microglia in FM group exhibited decrease in the number of intersections across the arbor in 3D Sholl analysis, smaller number of branches, shorter process length and elevated volume of CD68 as compared to HC group in (**a**-**d**) the ventral horn of the spinal cord lumbar segment (HC: n = 7 mice/65 cells, FM: n = 7 mice/76 cells for **a-c**; HC: n = 7 mice/43 cells, FM: n = 7 mice/ 36 cells for **d**); (**e**-**h**) the dorsal horn of spinal cord thoracic segment (HC: n = 7 mice/58 cells, FM: n = 7 mice/75 cells for **e**-**g**; HC: n = 7 mice/48 cells, FM: n = 7 mice/ 33 cells for **h**); (**i**-**l**) the ventral horn of spinal cord thoracic segment (HC: n = 7 mice/60 cells, FM: n = 7 mice/58 cells for **i**-**k**; HC: n = 7 mice/36 cells, FM: n = 7 mice/33 cells for **l**), and (**m**-**p**) anterior cingulate cortex (ACC) (HC: n = 7 mice/108 cells, FM: n = 7 mice/94 cells for **m**-**o**; HC: n = 7 mice/193 cells, FM: n = 7 mice/171 cells for **p**). Two-tailed, unpaired Student *t*-tests were used for statistical analysis in **b-d, f-h, j-l, n-p**. Two-way ANOVA followed by Tukey’s post-hoc comparison was used in **a**,**e**,**i**,**m**. \**P* < 0.05, \*\**P* < 0.01, \*\*\**P* < 0.001, \*\*\*\**P* < 0.0001. All data are presented as mean ± s.e.m.

**Extended Data Fig. 9.**
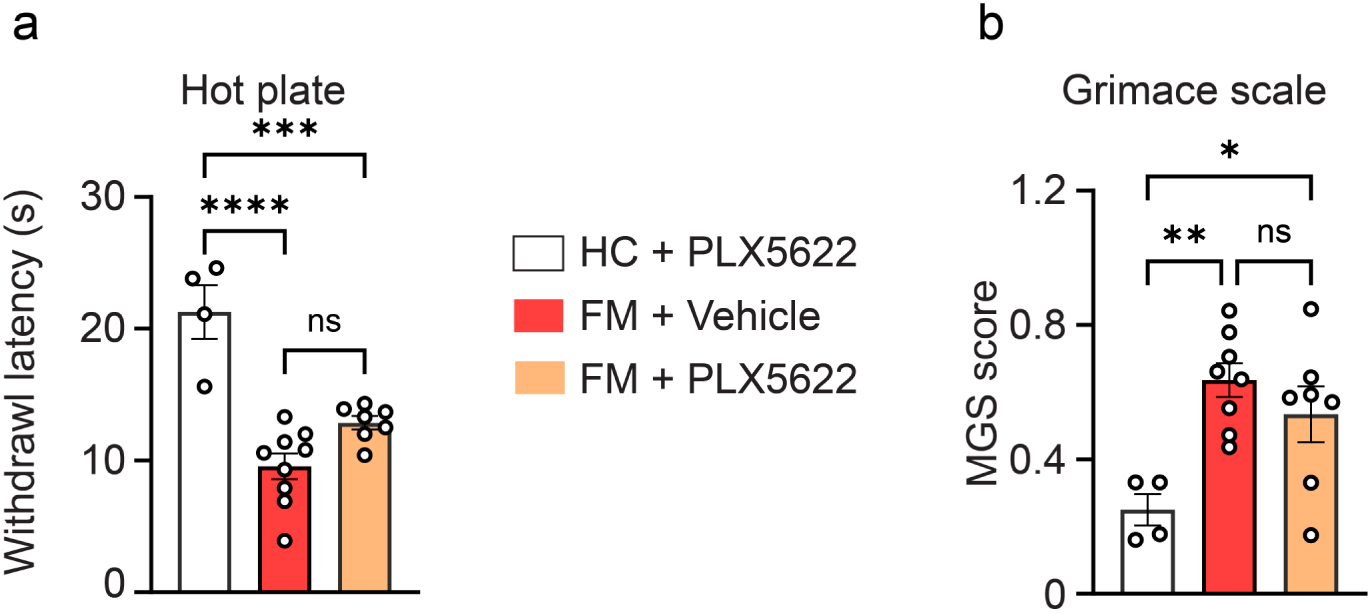
Effects of microglia depletion on heat sensitivity and spontaneous pain. Germ-free (GF) mice were treated with PLX5622 before receiving FMT from HC and individuals with FM. FM FMT in microglia-depleted mice in hot plate test (**a**) and spontaneous pain in the Mouse Grimace Scale (MGS) test (**b**). HC+PLX5622 (n = 4 mice), FM+Vehicle (n = 9 mice), FM+PLX5622 (n = 7 mice). One-way ANOVA followed by Tukey’s post-hoc comparison was used in both tests. \**P* < 0.05, \*\**P* < 0.01, \*\*\**P* < 0.001, \*\*\*\**P* < 0.0001. All data are presented as mean ± s.e.m.

**Extended Data Fig. 10.**
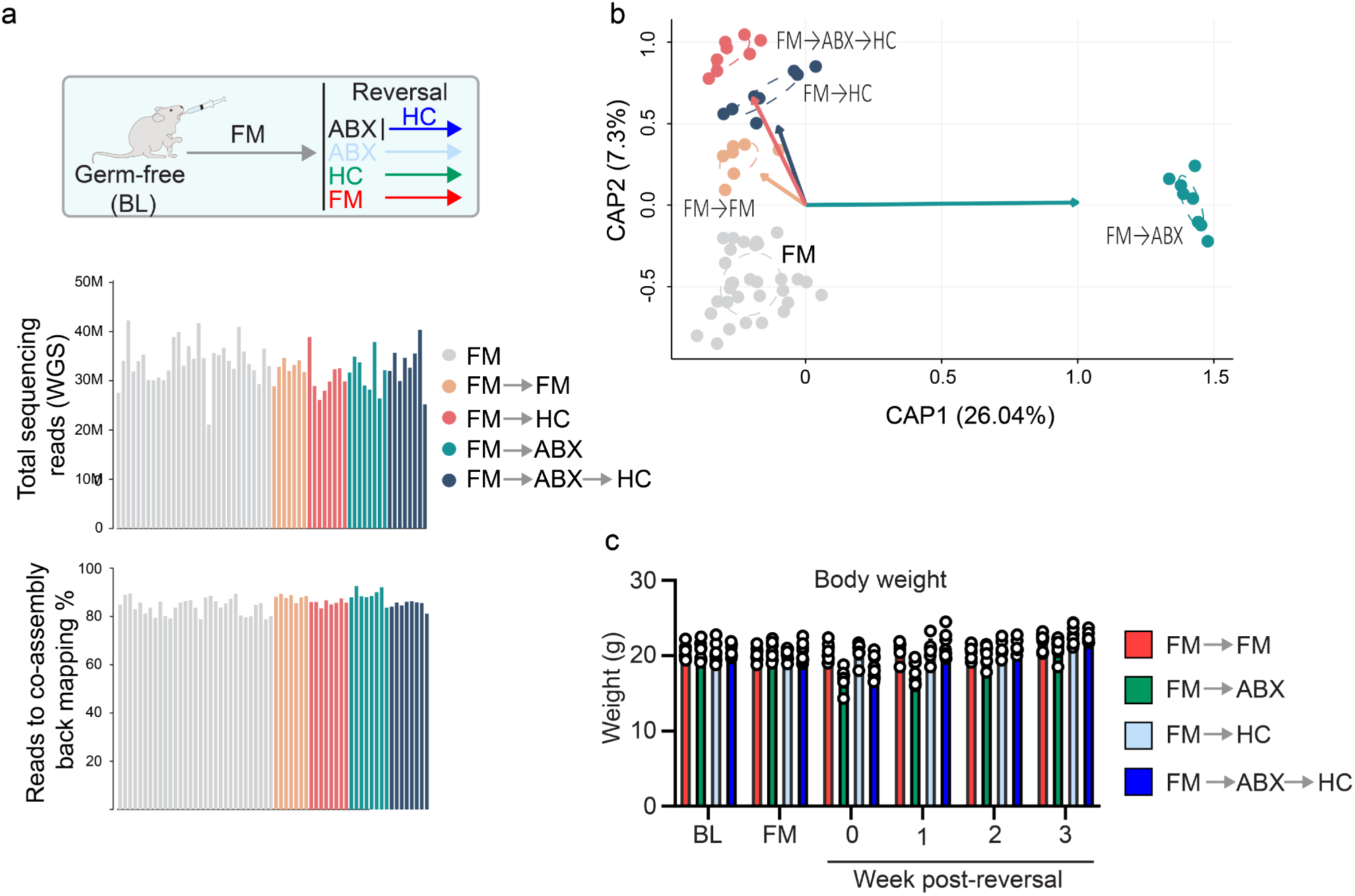
Whole genome sequencing shows differences in gut microbiota. **a,** Experimental design and sequencing characterization. Feces were collected at week 4 post-FM FMT and at week 4 post-reversal, and whole-genome sequencing was performed. Graphs in (**a)** show total sequencing reads (top graph) and percent of reads to co-assembly back mapping (bottom graph). **b**, CAP based on Bray-Curtis distances of normalized (log) WGS contig abundance of mouse samples at week 4 after FM FMT (FM) and at week 4 post-reversal (different interventions are labeled on the graph); The relative length of each vector indicates its contribution to each CAP axis shown. **c**, Assessment of body weight showed no differences between the groups. Data are presented as mean ± s.e.m. (n = 7-8 mice/group, also see Fig. 3).

**Extended Data Fig. 11.**
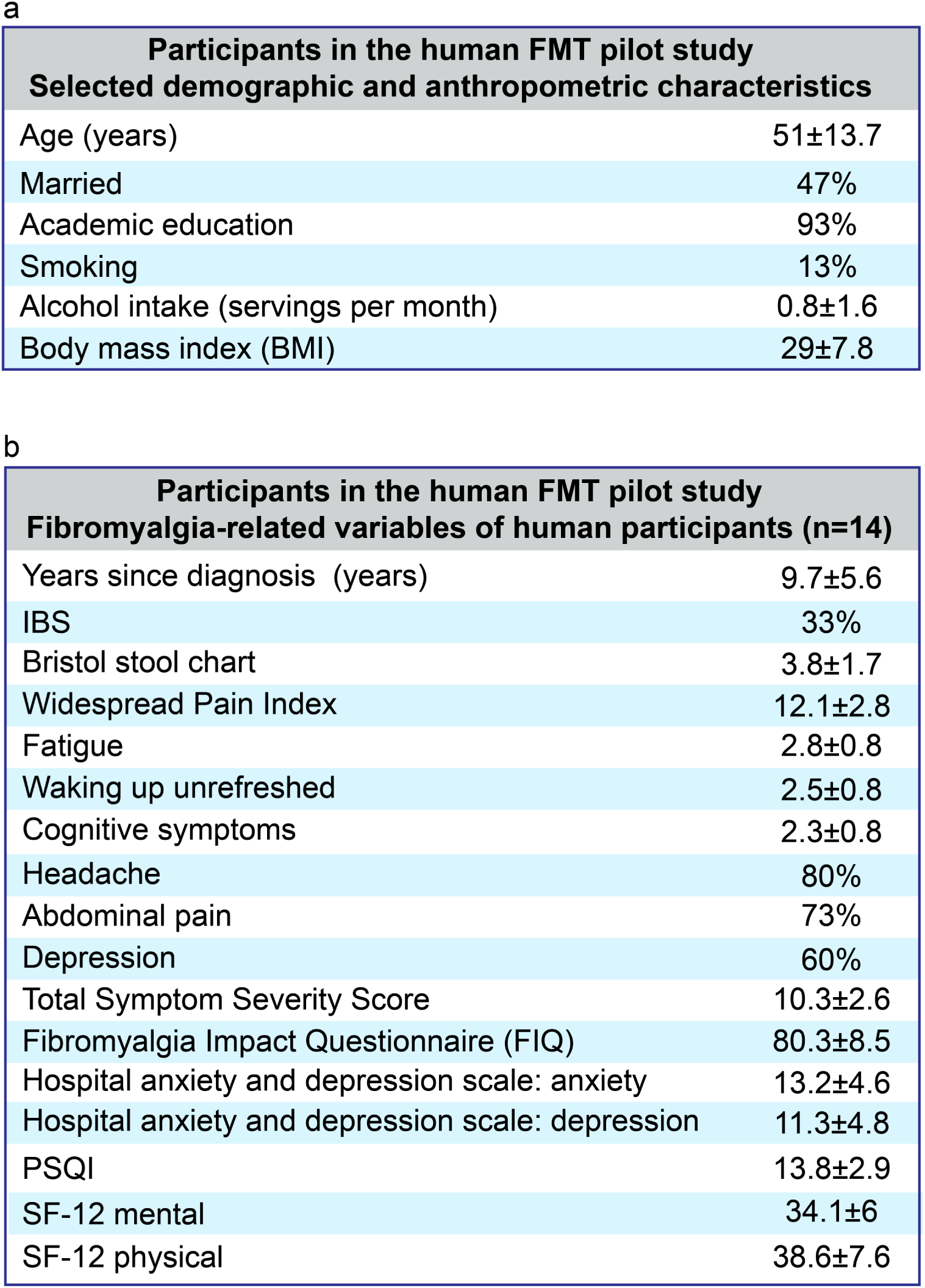
Characterization of participants in the human FMT study. **a,** Demographic and anthropometric characteristics of human participants in the FMT study (mean ± SD or % as indicated). **b**, Fibromyalgia-related variables of human participants: Years since diagnosis, co-diagnosis of IBS, disease severity and quality of life metrics based on the American College of Rheumatology Widespread Pain Index and Symptoms Severity Scale, Fibromyalgia Impact Questionnaire, and on the HADS, PSQI and SF-12 questionnaires (mean ± SD or % as indicated).

**Extended Data Fig. 12.**
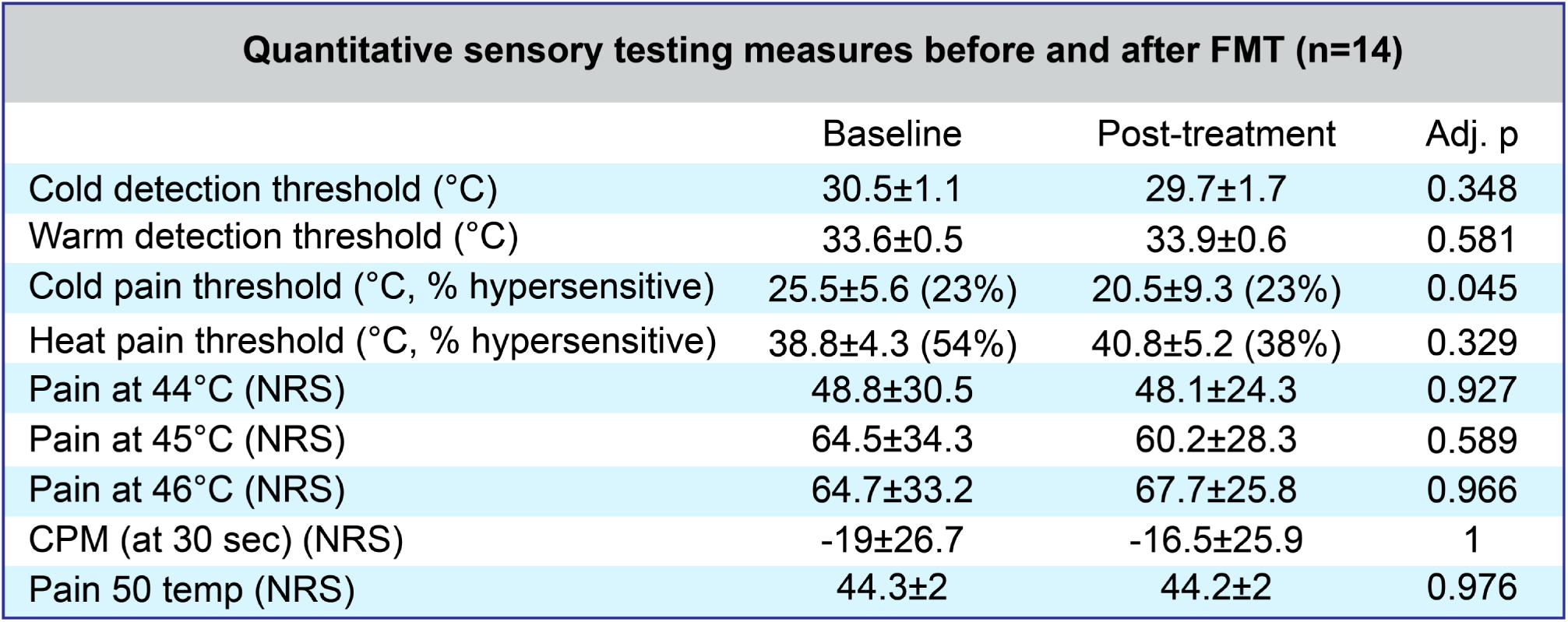
Quantitative sensory testing measures before and after FMT. Quantitative sensory testing measures before and after FMT: cold and warm detection thresholds; cold and heat pain thresholds including the proportion of patients whose thresholds fall beyond the normal ranges (see Methods); reported pain at 44, 45 and 46 °C; conditioned pain modulation (CPM) calculated by subtracting the test-stimulus pain ratings at 30 seconds from the test + conditioning pain ratings at 30 seconds; pain 50 was calculated as the average temperature at which patients reported pain intensity of 50 (NRS). Values are presented as mean ± SD. *P* values were adjusted using the Benjamini-Hochberg False Discovery Rate equation.

**Extended Data Fig. 13.**
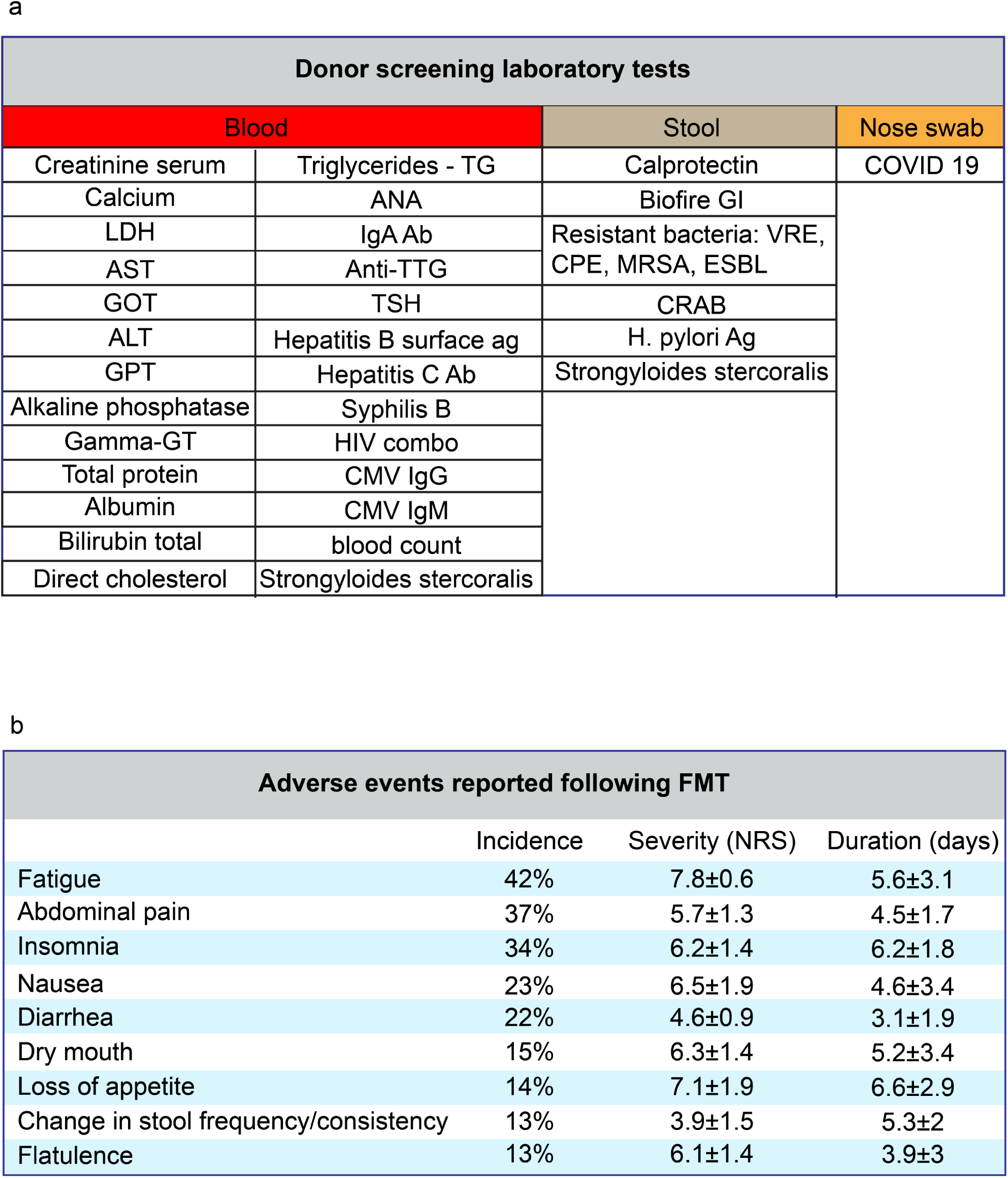
Characterization of donors in the human FMT study. **a,** Donor screening laboratory tests, including blood, stool and nose swab, which were taken in addition to a detailed intake questionnaire. Only candidates who screened negative for all questions and tests were eligible as stool donors. **b**, Adverse events (AE) reported following FMT, sorted by incidence. Incidence was evaluated as the mean number of patients reporting an AE following each FMT treatment divided by the number of participants. The average AE severity and duration are presented as mean ± SD. Post-FMT fatigue was the most common and severe AE, leading to the drop-out of 3 participants.

**Supplementary Table 1. Changes in the abundance of bacterial species at week 4 post-FMT in mice**

**Supplementary Table 2. Metabolomics changes in germ-free mice after transplantation of microbiota from fibromyalgia patients and healthy controls**

**Supplementary Table 3. scRNAseq of PBMCs in mice at week 4 post-FMT**

## Materials and Methods

### Animal study

#### Recruitment of human fecal donors

Recruitment of human participants took place at the Alan Edwards Pain Management Unit of the McGill University Health Centre, Montreal, Quebec, Canada. The study was approved by the McGill University Health Centre, Montreal institutional review board (2019-5521). All participants were given a detailed explanation of the study and signed an informed consent form. Participants were selected from the Fibromyalgia Gut Microbiome study cohort, which took place in the years 2017- 2019^1^. The cohort comprised 77 women with fibromyalgia and 79 healthy controls. Of these, women with fibromyalgia and healthy women were recruited as stool donors, by contacting individuals who had expressed their consent to be contacted for further studies. Donors were selected based on their clinical features (Extended Data Fig. 1a, b) as well as the similarity of their microbiome composition to the distribution of fibromyalgia and healthy control compositions as previously reported^2^. Briefly, inclusion criteria were: women; 30–60 years of age; a diagnosis of fibromyalgia (patient group) and no other acute or chronic medical condition (both groups); for fibromyalgia participants: Chronic Widespread Pain Index ≥9 and pain intensity ≥6 on a visual analogue scale; and ability to provide informed consent. Exclusion criteria were: any chronic illness, any acute illness in the preceding month, antibiotics treatment in the preceding 2 months, elevated anxiety or depression scores (HADS questionnaire >7 for anxiety or depression), irritable bowel syndrome (IBS) as per ROME IV criteria, serum C-reactive protein (CRP) levels >4 mg/L, fever >38 °C, pain scores >2 on a visual analogue scale (control group), pregnancy. Finally, representative microbiome compositions of fibromyalgia group were selected as follows: statistically significantly different multidimensional Mahalanobis distances between human patients (16S rRNA analysis of 77 samples of women with fibromyalgia ^2^; Extended Data Fig. 1c; see more below) and the respective distribution of fibromyalgia, or healthy gut bacterial compositions were calculated. Only samples with p-value higher than 0.1 (Chi-Square statistic of Mahalanobis distances) were selected as representative of the FM group.

Demographic and clinical variables were evaluated for normality of distribution using the Wilk-Shapiro test. Normally distributed variables were compared using paired-sample t-test, while non-normally distributed variables were compared using the non-parametric Wilcoxon signed-rank test.

Non-parametric repeated measure comparisons were done using the Friedman One-Way Repeated Measure Analysis of Variance. Adjustment for multiple comparisons was performed using Benjamini-Hochberg False Discovery Rate correction. Analyses were done on IBM_®_ SPSS® Statistics version 28.

#### Collection of stool samples and preparation of fecal microbiota transplants (FMT)

Fresh stool samples were collected by all participants at their homes, using a dedicated kit (Fisherbrand™ Commode Specimen Collection System), and delivered to the study facility within two hours of their collection. Samples were immediately processed, aliquoted, marked, and stored at -80 °C. A small volume of each sample was reserved for DNA extraction and further analysis using next-generation-sequencing. The remaining sample was stored in -80 °C.

For FMT sample preparation, an aliquot of the frozen sample was pulverized in a Biosafety class II hood with a ceramic mortar and pestle immersed in dry ice. An aliquot (1 g) of the pulverized material, sealed in a sterile screw-capped tube (Axygen SCT-200-C-S), was brought into an anaerobic chamber, immediately suspended in 15 ml PBS supplemented with 0.1% L-cysteine (in 50-ml conical shaped polypropylene tubes (Falcon)) and vortexed on maximum speed (four cycles of blending for 20 s followed by a 30-s pause). The sample was allowed to stand for 5 min so that particulate matter could settle by gravity, the resulting supernatant was passed through a 100-µm pore diameter filter to remove remaining particulate material, mixed with an equal volume of pre-reduced PBS (supplemented with 0.1 % L-cysteine) containing 30% glycerol (final concentration 15% glycerol). An aliquot (1.8 ml) of the suspension was placed in a 2.0 ml screw cap microtube and frozen at -80 °C.

#### Animals and environment

Female germ-free (GF) C57BL/6 mice (8-10 weeks of age) were purchased from Charles River (Canada). The animals were housed in flexible film gnotobiotic isolators (with surface sterilization achieved by treatment with Clidox) throughout the study, provided with autoclaved bedding, water, and food. Female C57BL/6J specific-pathogen-free (SPF) mice (8-10 weeks of age) were bred at McGill University from breeders originally obtained from The Jackson Laboratory (Bar Harbor, ME). All animals were maintained under a standard ambient temperature ranging from 21–23 ℃ and a light/dark cycle of 12:12 h (lights on at 07:00 h). Food (Envigo Teklad 8604) and water were provided *ad libitum*. All experiments were approved by the Downtown Animal Care Committee at McGill University and complied with Canadian Council on Animal Care guidelines (approval reference number FACC-B 7869). In all experiments, animals were randomly assigned to different groups.

Sample sizes were determined based on previous studies in the field. The experimenter was blinded to the condition in all studies.

#### Antibiotic treatment

Antibiotic treatment was applied as previously described^36^. Mice were provided autoclaved drinking water supplemented with a broad-spectrum antibiotic cocktail, containing ampicillin (0.5 mg/ml, Sigma), gentamicin (0.5 mg/ml, Sigma), metronidazole (0.5 mg/ml Sigma), neomycin (0.5 mg/ml, Sigma), vancomycin (0.25 mg/ml, Sigma), and saccharin (4 mg/ml, Sweet’N Low, Cumberland Packing Corp.). Sweet’N Low was added to make the antibiotic-containing water more palatable.

For antibiotics treatment model, specific pathogen-free (SPF) female mice were treated with a broad-spectrum antibiotic cocktail in drinking water for one week (as described above), followed by a one-time oral gavage of 200 microliters of macrogol, containing 12 mg of macrogol (polyethylene glycol, PEG) 4000 (colonsteril, Orion Pharma, Espoo, Finland). The mice then received weekly fecal microbiota transplantation (FMT) from healthy control (HC) or individuals with fibromyalgia (FM) for 5 weeks. For reversal experiment, the fibromyalgia FMT-recipient mice in FM-ABX-HC group were pre-treated with broad-spectrum antibiotic cocktail and PEG as above, and then received weekly FMT from HC.

#### Fecal microbiota transplantation into mice

For the fecal microbiota transplantation (FMT), recipient mice were administered a weekly dose of the supernatant solution by oral gavage at 10 ml/kg body weight, which equates to ∼ 200 µl per mouse. Tubes (2.0 ml screw cap microtube) were thawed and transferred into gnotobiotic isolators. To minimize unwanted exogenous colonization, both before and after FMT, the HC recipient mice and FM recipient mice were separately housed in two isolators. Within these isolators, they were kept in sterilized/autoclaved gnotobiotic iso-cages, 4-5 mice per cage. Behavioral testing started at baseline before FMT, while tissues were collected at the endpoint. Three healthy control (HC) and three fibromyalgia (FM) human donors were used in mouse transplantation experiments. The donors were carefully selected and characterized (Extended Data Fig. 1a-c). Each donor sample was transplanted to a group of 2-8 mice.

#### Behavioral studies

Behavioral studies with GF mice were performed inside the isolators, except radiant heat paw-withdrawal and cold-/hot-plate test, before and after the FMT. Experiments with SPF mice were performed in regular testing rooms. All experiments took place during the light cycle, no earlier than 09:00 h and no later than 16:00 h.

For the von Frey test, mice were placed in custom-made Plexiglas cubicles (5.3 × 8.5 × 3.6 cm) on an elevated wire mesh screen and habituated for 1 h. The 50% withdrawal thresholds were estimated using von Frey filaments (calibrated, Stoelting Co, Wood Dale, IL) by the “up-down” method of Dixon^37^. Filaments were firmly applied to the plantar surface of both hind paws for 3 sec and a rapid paw withdrawal indicates a positive response. Mice were tested for mechanical sensitivity at baseline and weekly until the end time points after fecal microbiome transplantation to assess mechanical thresholds.

For the radiant heat paw-withdrawal (Hargreaves’) test, mice were habituated in custom-made Plexiglas cubicles (5.3 × 8.5 × 3.6 cm) on a glass plate for 1 h prior to testing. A beam of light that was emitted from the light box was applied to the middle of the plantar surface of each hind paw to stimulate. The light beam was set at 20% of the maximum (IITC Model 390) and was turned off when a rapid lift or withdrawal of the hind paw occurred. We made 2∼3 consecutive measures on each hind paw at each time point that were then averaged. A cut-off time of 30 sec was applied to avoid tissue damage.

For the cold-/hot-plate test, the mouse was placed in a Plexiglas cylindrical testing chamber on a hot plate set at 52.8 °C or a cold plate set at 3.5 °C (Ugo Basile srl, Model 35150). The latency to react to the hot or cold stimuli was quantified as the length of time between the placement of the mouse on the plate and the occurrence of licking/lifting of the paw(s) or jumping. A cut-off time of 30 sec was applied to avoid tissue damage.

The Mouse Grimace Scale (MGS) test was modified slightly from the original description^38^. Mice were placed in custom-made Plexiglas cubicles (5.3×8.5×3.6 cm) on a perforated metal floor and were habituated for 30 min before testing. Mice were recorded for 1 h with a digital video camera (Sony Digital Camcorder HDR-PJ430V). A total of 20 head-shot photos were used to define a mean score for each animal, in which one photo was taken for every 3-min interval within the 1-h recording. Photos of sleeping mice were discarded as well as photos of mice expressing grooming or scratching behavior. Coders were subsequently blinded, and photos were randomized and then scored using the following criteria: the intensity rated as a value of 0, 1 or 2 for each of the five action units (AUs), which include orbital tightening, nose bulge, check bulge, ear position and whisker change. In every case, 0 indicated AU was not present, 1 indicated moderate visibility of the AU, and 2 indicated severe changes in the AU. An MGS score for each photo was calculated by averaging intensity ratings for each AU.

The elevated plus maze (EPM) consists of four Plexiglas arms: two closed and two open, positioned 50 cm above the ground (model H10-35-EPM). A slightly raised lip, 0.5 cm in height, surrounds the perimeter of the open arms, ensuring the mouse does not fall. Each mouse was placed individually at the center of the maze, facing an open arm, and allowed to explore freely for 5 min under overhead fluorescent lighting (200 lux). A digital video camera (Sony Digital Camcorder HDR-PJ430V) was situated above the maze to capture a top-down perspective. The time each mouse spent in different arms was recorded, and the percentage of the total 5 minutes was compared between groups.

The novel object location (NOL) test spanned four consecutive days with three distinct phases. Day one involved a 10-min habituation in an empty 60 cm x 60 cm x 30 cm box. The next two days had two 10-min sessions where mice explored the box with two identical objects placed near opposite walls. On day four, during a 10-min test, one object was moved to a corner. Mice activity was recorded using a Sony Digital Camcorder HDR-PJ430V. A discrimination index was calculated by comparing the time spent exploring each object relative to the total exploration time.

For the tail-suspension (TST) test, mice were suspended by their tails and secured with PVC white vinyl insulation tape for 6 min. To prevent mice from climbing up their tails during the test, hollow cylinders made from cut 3-mL syringes were placed around the tails. A digital recording camera was used to record. A stopwatch was used to measure the duration of each mouse’s immobility versus escape attempts, which included body shaking, running-like motions, and climbing attempts.

#### Peripheral blood mononuclear cell (PBMCs) collection

Blood (200 µl) was collected from the cheek vein into EDTA-coated tubes, diluted 1:3 with a balanced salt solution (1X Dulbecco′s PBS (DPBS, Cat No. TMS-012-A Sigma Aldrich) fortified with 2% FBS (Cat No. F2442, Sigma Aldrich)), and transferred to 15 ml SepMate™ PBMC isolation tubes (Cat No. 85415, STEMCELL Technologies). A 4.5 ml Ficoll^®^ Paque Plus solution (Cat No. GE17-1440-02, Sigma Aldrich) was added beneath the diluted blood, and tubes were centrifuged at 400 x *g*. The buffy coat layer was harvested and rinsed with the balanced salt solution (3X the volume of the buffy coat layer). Subsequently, it was centrifuged at 400 x *g* for 10 min at room temperature.

For platelet removal, we adapted the method from ref. ^39^. Briefly, the cell pellets were resuspended in 1 ml DPBS and a sucrose gradient is created in a 15-mL tube with 4 mL of 10% sucrose and 2 mL of 15% sucrose with resuspended cells carefully layered on top. After centrifugation at 200 x *g* for 15 min (no acceleration and no break set on the centrifuge), the top 5 mL of supernatant is discarded, and the remainder is centrifuged at 500 x *g* for 5 mins to get cell pellet. Add 1 ml ACK lysing buffer (Cat No. A1049201, Thermo Fisher Scientific), incubate at room temperature for 5 mins, centrifuge at 300 x *g* for 5 mins at room temperature. Aspirate the supernatant, leaving approximately 50 µl to avoid disturbing the pellet. Gently mix the cells and remaining fluid, then wash the cells by adding 5ml balanced salt solution and centrifuging at 500 x g for 10 min at room temperature. Finally, cells are resuspended in 100 µl of 0.04% BSA/DPBS and are ready for scRNA sequencing.

#### Single-cell RNA sequencing and analysis

Approximately 9,200 cells from each sample were processed using a Chromium Single Cell Chip (10x Genomics) and the Chromium Single Cell 3ʹ Library & Gel Bead Kit v3 (10X Genomics) for reverse transcription and library preparation. Sequencing, averaging 67,000 reads per cell, was done on an Illumina NovaSeq sequencer. Sample demultiplexing, barcode processing, gene counting, and mapping to the reference transcriptome (mouse mm10 v.1.2.0) were handled by Cell Ranger (v.3.0.1) (10X Genomics).

Using the Cell Ranger software provided by 10X Genomics (version 3.0.2), the raw sequencing data from Illumina HiSeq’s BCL files was converted to expression counts matrices. The data were demultiplexed into gzip-compressed FASTQ files and aligned to the GRCm38 (mm10) mouse reference genome. UMI count matrices (genes × cells) were then fed into Seurat (version 4.1.0)^40^. Cells with gene counts outside the range of 200–2,500 or with over 10% of transcript counts from mitochondrial genes were excluded and the doublets detected by the DoubletFinder R package were filtered out^41^. Following data normalization, feature extraction, and dataset scaling, we conducted principal component analysis (PCA) and uniform manifold approximation and projection (UMAP). A resolution of 0.2 was used to categorize the cells into 12 distinct clusters using the FindClusters function. To earmark cluster-specific markers, we adopted analogous thresholds with the FindAllMarkers function. Cluster-specific features were applied to annotate cell types based on known canonical markers: memory B cells (Cd79a^+^Cd19^+^), plasma B cells (Cd79a^+^Cd19^-^), B cell-like T cells (Cd79a^+^Cd19^+^Cd4^+^), classical monocyte clusters (Cd44^+^Ly6C2^high^), intermediate monocyte cluster (Cd44^+^Ly6C2^int^), non-classical monocyte clusters (Cd44^+^Ly6C2^low^), CD4 naïve T cells (Cd4^+^Ccr7^+^Cd3d^+^), CD8 naïve T cells (Ccr7^+^Cd3d^+^Cd8a^+^), T4_Treg cells (Cd3e^+^Cd4^+^Cd24a^+^), NK cells (Ncr1^+^Gzma^+^), DC (Itgax^+^Fcer1a^+^), platelet (Pf4^+^Ppbp^+^). We determined differentially expressed genes (DEGs) by establishing cutoffs: a |log fold-change| exceeding 0.25 and a Benjamini-Hochberg FDR below 0.05. Gene ontology analyses were conducted using Enrichr^42^.

#### Microbial community analysis

Microbial communities from stool samples of human donors and of mice receiving FMT from HC or FM human donors were assessed using 16S rRNA gene sequencing and whole genome sequencing (WGS) accounting for 406 and 90 samples, respectively. The ANCHOR pipeline^43^ was used to process amplicon sequences and infer exact sequence variants (ESVs) while WGS samples were co-assembled, contigs were annotated and functionality was explored by predicting genes on assembled contigs. Briefly:

#### 16S rRNA gene and WGS assemblies

DNA from 406 samples was extracted at Genome Quebec laboratories and V5V6 region of the 16S rRNA gene was amplified using P609D-GGMTTAGATACCCBDGTA (forward) and P699R-GGGTYKCGCTCGTTR (reverse) primer set. A total of 26,199,758 amplicons were sequenced. Samples were analyzed using 3 separate instances of Anchor pipeline. The first one, labeled FM FMT contained 144 samples, the second, FM intervention contained 186 samples, and the third, Donor vs FMT mice contained 76 samples.

Microbial communities from stool samples of human donors and of mice receiving FMT from human donors with and without FM were assessed using 16S rRNA gene sequencing, and the ANCHOR pipeline was used to process amplicon sequences and infer exact sequence variants (ESVs). Mothur ^44^ was used to assemble reads into contigs and for dereplication. Anchor threshold was set to 42 (FM FMT) and 20 (FM reversal). Two databases (NCBI nt and NCBI 16S curated database; January 2022 versions) were used for annotation using BLASTn (v 2.10.0+;^45^) as the alignment search tool with the following parameters: a minimum of 99% identity and coverage for taxonomy inference. USEARCH (v9;^46^) was used to flag chimera sequences.

DNA from 90 samples was extracted and sequenced at Genome Quebec laboratories on Illumina NovaSeq 6000 S4 PE150 sequencing machines using whole genome sequencing protocol. Trim Galore!^47^, a wrapper based on cutadapt^48^ and fastqc^49^, was used to perform a quality control step on the raw paired-end reads with the following parameters: --trim-n --max_n 0 --paired --retain_unpaired --phred33 --length 75 -q 5 --stringency 1 -e 0.1 -j 1. BBMAP ^50^ was used to remove potential contamination from human using the masked version of hg19 human assembly. BBMAP was also used to remove mouse DNA using mm10 mouse genome. Ribosomal RNA was removed using SortmeRNA^51^. MEGAHIT^52^ was used to assemble reads from all samples into one co-assembly using meta-large option. Kallisto^53^ expectation maximization algorithm was used to complete metagenomics read assignment and infer contig abundance^54^. To assign contig taxonomy, alignment was run using full contig lengths against the NCBI nr/nt database (January 2022) and Reference Viral Database (RVDB v v25.0). BLASTn was run using the following parameters: -evalue 1e-50 - word_size 128 -perc_identity 97.

#### Microbiome data analysis

Sparsity filter was set to 0.9 and minimum count occurrence per analyzed group was set to 3 samples. DESeq2^55^, Phyloseq^56^, Vegan^57^ and ggplot2^58^ R libraries were used to do alpha (based on Shannon index) and beta (bray-curtis distance based unsupervised and supervised ordinations) diversity analysis and differential abundance (DA) analysis. Correlation analysis was produced from corrplot^59^ and Psych^60^ R libraries using Kendall rank correlation. Correlation and ordination diagrams were based on regularized-logarithm transformation of raw counts (rlog function; DESeq2). Flower diagram was made using custom JavaScript script. Significance on alpha diversity was obtained using either parametric (t-test) or non-parametric (Mann–Whitney U test) depending on the index value distribution. Significance on beta diversity was evaluated with Monte Carlo permutation testing between sample groups (10,000 permutations). Multiple t-tests were applied to the individual correlations and p-values were adjusted for multiple tests using Benjamini-Hochberg procedure. In DA analysis the significance of the adjusted p-value was set for values lower or equal to 0.1. To control for technical replication, we applied a series of conditions: 1) only differentially abundant ESVs in mice originating from all donors were retained; 2) Sparsity (values representing, for a given ESV, the total count range among samples; 1 representing a total count originating from a single sample), calculated from the average of all mice from a same donor, was evaluated and only ESVs with a sparsity lower of 0.9 were retained; 3) Only ESV with low variance between technical replicates (coefficient of variation <0.1) were retained. Coefficient of variation was calculated as is the ratio of the standard deviation to the mean between average counts of the samples from the same group (FM or HC).

#### Selected FM sample representativeness

Three samples were selected as a subset of the initial FM group (77 samples in total^2^). Multivariate representation (PCoA ordination) was chosen as the method to address subs selection’s FM group representativeness. PCoA ordination based on Bray-Curtis distance was built based on regularized logarithm transformation of 16S rRNA ESV raw abundance (rlog function from DESeq2 R library) and potential outliers were excluded using the Mahalanobis distance (i.e. distance between each sample and the ordination distribution). Sample Mahalanobis distances (mahalanobis function in R) were estimated using PCoA’s principal component scores (5 first principal components), the mean vector of the distribution and the variance-covariance matrix of the distribution. To identify Mahalanobis distances that were statistically significantly different from the FM group distribution (i.e. larger distances representing samples that are "far" from the group distribution), p-values were calculated using the Chi-Square statistic of Mahalanobis distances (pchisq function from R). Only samples with p-value higher than 0.1 were selected as representative of the FM group.

#### Immunohistochemistry and imaging

Mice were anesthetized and perfused intracardially using PBS, followed by 4% paraformaldehyde (PFA) in 0.1 M phosphate buffer (pH 7.4). The brain, spinal cord and glabrous hind paw skin were then extracted and post-fixed in 4% PFA in phosphate buffer at 4 °C for 24 h. Following post-fixation, spinal cords and brains were transferred to PBS, sectioned at 30 μm for microglia measurement, or 70-μm thickness for microglia morphology study using a vibratome, and collected as free-floating in PBS in 24-well plates. Skin tissues were immersed 24 hours post-transfer in a cryoprotectant solution composed of 30% sucrose in 0.1M PB. They were then embedded in OCT (Thermo Fisher Scientific), sectioned at 14 μm using a Leica cryostat, and collected directly onto gelatin-subbed histological slides.

For microglia measurement, free-floating sections (30 μm) were blocked with 10% normal donkey and goat serum (NGS/NDS) in 0.2% Triton-X in PBS (PBS-T) for 1 h, then incubated with primary antibodies (Iba-1 ([1:500], Wako 019-19741), and CD68 ([1:500] Bio-Rad MCA1957)) diluted in a 5% NDS/NGS solution PBS-T at 4 °C for 24 h. Subsequently, sections were washed three times in PBS-T, each for 10 min, and incubated with secondary antibodies (1:500 Goat anti -rabbit Alexa Fluor 647 [Thermo Fisher Scientific, A-21245], 1:500 Donkey anti-rat Alexa Fluor 594 [Thermo Fisher Scientific, A-21209]) in PBS for 2 h at room temperature. Following three additional 10-min washes in PBS, tissues were mounted using ProLong™ Gold Antifade Mountant with DNA Stain DAPI (Thermo Fisher Scientific, P36931).

For microglia measurement study, Z-stacks were taken at 20X magnification, Airyscan processed, and processed as maximum intensity projections of confocal z stacks using ImageJ software (v.1.52). Each group consisted of a total of n = 4 mice, with 2-3 sections of spinal cord being taken from each mouse.

For microglia morphology study, free-floating sections (70 μm) were blocked with 10% normal donkey and goat serum (NGS/NDS) in 0.2% Triton-X in PBS (PBS-T) for 2 h, then incubated with primary antibodies (Iba-1 ([1:500], Wako 019-19741), and CD68 ([1:500] Bio-Rad MCA1957)) diluted in a 5% NDS/NGS solution PBS-T at 4 °C for 24 h. Subsequently, sections were washed five times in PBS-T, each for 10 min, and incubated with secondary antibodies (1:500 Goat anti -rabbit Alexa Fluor 647 [Thermo Fisher Scientific, A-21245], 1:500 Donkey anti-rat Alexa Fluor 594 [Thermo Fisher Scientific, A-21209]) in PBS for 3 h at room temperature. Following five additional 10-min washes in PBS, tissues were mounted using ProLong™ Gold Antifade Mountant with DNA Stain DAPI (Thermo Fisher Scientific, P36931).

For microglia morphology study, Z-stacks were taken using 63x/1.4 Oil DIC M27 objective, Airyscan processed, and processed as maximum intensity projections of confocal z stacks using ImageJ software (v.1.52). Each group consisted of a total of n = 7 mice, with 4 sections of brain and spinal cord being taken from each mouse.

For intraepidermal nerve fiber (IENF) density assessment in the glabrous skin, skin section slides were washed with PBS to remove residual OCT. Section slides were blocked with 10% NGS/NDS in PBS-T for 1 h, then incubated with primary antibodies PGP9.5 ([1:400], PA5-29012, Invitrogen) diluted in a 5% NDS/NGS solution PBS-T at 4 °C for 48 h. Subsequently, section slides were washed three times in PBS-T, each for 15 min, and incubated with secondary antibody (1:500, Goat anti - rabbit Alexa Fluor 568, Thermo Fisher Scientific, A-11011) in PBS for 2 h at room temperature. Following three additional 10-min washes in PBS, section slides were mounted using ProLong™ Gold Antifade Mountant with DNA Stain DAPI [Thermo Fisher Scientific, P36931].

For IENF study, imaging was performed in Airyscan mode using a Zeiss Apochromat 63x/1.4 Oil DIC M27 objective along with the Airyscan super-resolution (SR) module, equipped with 32-channel hexagonal array GaAsP detector on LSM880 (Zeiss) using a 561-nm laser. Airyscan super-resolution image stacks were reconstructed using ZEN Black software (Zeiss). Each group consisted of a total of n = 6 mice, with 5-6 sections taken from each mouse.

#### Image analysis quantification

The quantification of sensory fibers in the paw skin and cornea was performed with ImageJ software. Prior to any type of analysis, a consistent live frame area was measured (100 μm × 100 μm) and placed in an area where fibers were most dense. The intraepidermal nerve fiber counting rule was adapted from ref. ^61^. Nerve fibers which cross the basement membrane were counted as one nerve fiber. Nerve fiber which branches after crossing the basement membrane or which resides only in the epidermis was excluded. The epidermal nerve fiber branches before crossing the basement membrane were counted as two fibers.

The Sholl analysis of microglia in the spinal cord was done using IMARIS (software version 9.7.2). Thick sections (70 μm) of the brain and spinal cord were used in IHC to amplify the Iba1 signal. High-resolution z-stacks of single microglia were taken using a Zeiss LSM700 upright confocal microscope using a 63X oil objective (NA: 1.4) (38-48 μm z-stacks; xy resolution: 0.10 μm; z step: 0.2 μm, 1024×1024 px). Imaris 9.7.1 file converter was used to change the CZI file type (.czi) to an Imaris file type (.ims) for access in Imaris 9.7.1. The images contained Iba1 and CD68 channels. In Imaris, the Filament Tracer Module was used to reconstruct the microglia cells. The Iba1 channel was selected as the primary channel for Iba1 microglia activation analysis. The other channel was deselected prior to filament reconstruction. The data type of interest included the “filament number of Sholl intersections.” Imaris produced an Excel data sheet with raw data, which includes: 1. Filament length, 2. Sholl analysis, 3. Total number of dendrites. Microglia morphology alteration was compared using quantitative data obtained from the following four locations in the spinal cord: lumbar dorsal horn, lumbar ventral horn, thoracic dorsal horn, thoracic ventral horn, and anterior cingulate cortex (ACC) of the brain.

IMARIS 3D reconstruction was used alongside representative images of Iba1/CD68 to show CD68 localization inside microglia.

#### PLX5622

PLX5622 administration was used based on the protocol previously described^62^. PLX5622 (Cat.# HY-113153/CS-0077157, MedChemExpress) was dissolved in 10% DMSO, 20% RH40 (KolliphorR RH40, Sigma, Lot# BCCD2285), 5% Tween-80, 65% saline (strictly in this order), making a working solution at 6.5 mg/ml. Mice received i.p injection daily dose at 65 mg/kg body weight for 7 days.

#### RNA Extraction and cDNA synthesis

Total RNA was extracted from de-identified tubes of mouse tissues using the RNeasy Kit (QIAGEN; Germany), according to the manufacturer’s instructions. RNA concentrations and the 260/280 nm absorbance ratio were determined using NanoDrop One (Thermo Fisher Scientific). cDNA was reverse transcribed using iScript cDNA Synthesis Kit (Bio-Rad; Hercules, CA, USA), according to the manufacturer’s protocol using 1 µg total RNA in 20-µL reactions.

#### Quantitative Real-Time PCR

qRT-PCR reactions were carried out in a total reaction volume of 10 µL containing: 5 µL of 2X Power SYBR Green PCR Master Mix (Applied Bioscience; Foster City, CA, USA), 0.4 µL each of 10-mM primers, 3.2 µL of double distilled water, and 1 µL of cDNA. qRT-PCR were performed in triplicates using a 96-well plate format in the ABI PRISM 7900 HT (Thermo Fisher Scientific) with the following conditions: 2 min at 50 °C and 10 min at 95 °C, followed by 40 cycles of 15 s at 95 °C and 1 min at 60 °C, dissociation stage consisted of 15 s at 95 °C, 15 s at 60 °C, 15 s at 95 °C. β-actin was used as house-keeping gene to normalize expression levels between samples. The following primers were used: *Actb*: Forward:5’-GGCTGTATTCCCCTCCATCG-3’, Reverse: 5’- CCAGTTGGTAACAATGCCATGT-3’; *Zo1*:Forward:5’-AGGACACCAAAGCATGTGAG- 3’,Reverse:5’-GGCATTCCTGCTGGTTACA-3’; *Zo2*: Forward: 5’- ATGGGAGCAGTACACCGTGA-3’, Reverse:5’-TGACCACCCTGTCATTTTCTTG-3’; *Ocln*: Forward:5’-TTGAAAGTCCACCTCCTTACAGA-3’,Reverse:5’- CCGGATAAAAAGAGTACGCTGG-3’.

#### Intestinal permeability assay

Four weeks after FMT, mice were fasted (food removal only) for 4 hours. Subsequently, mice were treated with 4 kDa FITC-dextran (100 mg/ml; Sigma; St. Louis, MO, USA), reconstituted to 80 mg/ml in PBS, administered at 600 mg/kg body weight by oral gavage. Four hours after gavage, mouse blood was collected by cheek puncture into 1.1-ml z-gel serum collection tubes (Sarstedt; Germany) and stored on ice in the dark at 4 °C. After a 30-min clotting period, samples were centrifuged at 10,000 x *g* for 10 min, and serum was transferred to a clean tube. Standard and samples were prepared in a fluorescence-compatible 96-well plate (Costar #3915) with 100 µl total volume per well. A standard curve was generated with serial dilutions and serum was diluted with an equal volume of 1X PBS with duplication. Fluorescence measured at 530 nm (excitation at 485 nm).

#### Metabolomics data analysis

Tissues were collected (brain, spinal cord, plasma, and feces) from mice at week 4 post-FMT, stored at 80 °C, and shipped to the Metabolon Inc. (Durham, NC, USA) for metabolomics analysis. All methods utilized a Waters ACQUITY ultra-performance liquid chromatography (UPLC) and a Thermo Scientific Q-Exactive high resolution/accurate mass spectrometer interfaced with a heated electrospray ionization (HESI-II) source and Orbitrap mass analyzer operated at 35,000 mass resolution.

#### Gut histology

Mice were sacrificed 4 weeks after FMT. The small intestine, cecum and colon were collected, opened longitudinally, and washed thoroughly with PBS then processed as a ‘Swiss roll’ in paraffin. Four-micrometer sections were cut and stained with hematoxylin and eosin [H&E] using a standard protocol. Sections were scored based on the existing literature^63^, and eight histological parameters: inflammatory infiltrate, goblet cell loss, hyperplasia, crypt density, muscle thickness, submucosal infiltration, ulcerations, and crypt abscesses, each scored from 0–3. A total histological severity score, ranging from 0–24, was calculated by summing the eight assessed parameters.

#### Statistical analysis for work involving mice

Animals were distributed into various treatment groups randomly. Appropriate sample sizes were estimated based on prior experiences in performing similar types of behavioral and molecular analyses. All data were expressed as mean ± s.e.m. The data were statistically analyzed with two-tailed, unpaired Student’s *t*-test and a One-way or Two-way ANOVA followed by Tukey’s post-hoc comparison test (GraphPad Prism 9.1.0), as appropriate. Significance was set at α = 0.05.

### Open label clinical trial

#### Ethics approval and oversight

The human study was conducted at the Institute for Pain Medicine of the Rambam Health Campus, Haifa, Israel, between October 2022 and October 2023. The study was approved by the Rambam Institutional Review Board (approval number RMB-0481-21). Participants and donors were given a detailed explanation of the study and signed an informed consent form.

#### Donors screening and capsulized FMT preparation

Healthy women, aged 18–60 years were screened as stool donors, based on the Israeli Ministry of Health regulations for stool donors^1^ (Extended Data Fig. 13a) and international guidelines^64^. Eligible donors delivered fresh stool samples in dedicated collection containers (Thermo Fisher Scientific, Massachusetts USA) and delivered on ice within 4–8 h of passing. Stool samples were processed within 1 h of their delivery, under strict anaerobic conditions in an anaerobic chamber (Bactron 300, Sheldon Manufacturing, Oregon USA) using an anaerobic gas mixture (90% N_2_, 5% CO_2_, and 5% H_2_). Samples were homogenized and grossly filtered in a laboratory blender (Bag Mixer, Interscience, France)^65^ as follows: Raw stool was transferred into a stomacher bag (BA6141, Seward, NY USA) containing sterile 20% glycerol in normal saline solution in a 1:1 ratio to a final glycerol concentration of 10%, and then homogenized using a stomacher machine (BagMixer 400 P, Interscience, France). The resulting slurry was double encapsulated in HydroxyPropylMethylCellulose capsules (Capsugel, Lonza, Switzerland). FMT capsules were separated into packages containing 15 capsules and immediately flash frozen in liquid nitrogen, to be then transferred to -80 °C for long-term storage.

#### Patient selection and baseline evaluation

Women with severe fibromyalgia resistant to treatment were recruited by advertisement in national fibromyalgia support groups. Inclusion criteria were: women aged 18–65 years diagnosed with fibromyalgia; average reported pain of ≥6 during the preceding week; Fibromyalgia Impact Questionnaire (FIQ) score of ≥60/100^66^; symptomatic despite receiving standard care for fibromyalgia. Exclusion criteria included active inflammatory diseases (e.g., rheumatic, gastrointestinal); malignant neoplasm in the preceding 5 years; immunosuppression; uncontrolled psychiatric pathology; known allergy to the planned antibiotics or to laxatives used in the preparation protocol; severe food allergy. Eligible participants were interviewed by a specialized pain physician for a thorough assessment. Only individuals whose diagnosis of fibromyalgia was confirmed, based on the 2016 Fibromyalgia Diagnostic Criteria^67^ were deemed eligible to participate in the study. Participants were interviewed and data were collected including demographics, anthropometrics, co-morbidities (including a specific evaluation for irritable bowel syndrome, based on ROME IV criteria^68,69^), medications, smoking, and alcohol consumption. Participants then filled-in the following questionnaires: The Fibromyalgia Survey Diagnostic Criteria and Severity Scale (FSDC) questionnaire, assessing symptom severity, pain location, fatigue, sleep quality and cognitive and somatic complaints in fibromyalgia patients, based on the 2016 criteria for the diagnosis of fibromyalgia^67^; The Fibromyalgia Impact Questionnaire (FIQ), a 10-item questionnaire evaluating physical functioning, work difficulty, pain, fatigue, morning tiredness, stiffness, anxiety and depression^70^; Hospital Anxiety and Depression Scale, a 14-item questionnaire aimed to measure anxiety and depression in a general medical population of patients^71^; the Pittsburgh Sleep Quality Index (PSQI) – a 19-item questionnaire assessing sleep quality over a 1-month time interval^72^; Short-form health-questionnaire (SF-12), a 12-item questionnaire evaluating the impact of health on an individual’s quality of life^73^; the Bristol stool form scale (BSFS), a one-item questionnaire designed to assess the form of human stool into one of 7 categories^74^.

#### Quantitative sensory testing (QST)

QST measurements were done at baseline, and one week after treatment termination.

All patients underwent quantitative sensory testing, including thermal detection and pain thresholds, thermal pain sensitivity evaluation (pain 50), and conditioned pain modulation (TSA-2001, Medoc, Israel and Heto CBN 8-30, Allerod, Denmark).

The method of limits was used to determine the thermal detection and pain thresholds^75^. Cool and warm detection thresholds (CDT and WDT, respectively) and heat and cold pain thresholds (HPT and CPT) were measured using a 30×30mm thermode of the Thermal Sensory Analyzer (TSA-II, Medoc Ltd, Ramat Yishay, Israel) system on the volar aspect of participants’ non-dominant hand. Each test included a series of five stimuli separated by a 4–6 second inter-stimulus interval (ISI).

For the evaluation of CDT and WDT, thermode temperature was decreased or increased at a rate of 0.5 °C/s from the baseline temperature of 32 °C. Participants were asked to press a button when they first sensed a cold or warm sensation. The thermode temperature recovery to baseline was set to a rate of 8 °C/s. The thermal detection threshold was calculated as the mean temperature of the three closest readings with an interval <0.5 °C.

For the HPT and CPT, thermode temperature was decreased or increased from baseline at a rate of 1 °C/s. Participants were asked to press the button when they first sensed pain. The thermode was slightly repositioned after each stimulus to avoid adaptation or sensitization. HPT and CPT were calculated as the mean of the three closest temperature readings, preferably with a difference <0.5 °C.

Individual Z-scores were calculated for the thermal sensory and pain thresholds and adapted for sex and age. Thresholds were based on the German Research Network on Neuropathic Pain DFNS reference^76^ (z-score = [mean participant threshold – mean reference threshold]/standard deviation). A z-score > +1.96 or < -1.96 was considered as pathological hypo-or hypersensitivity. Cold and warm detection thresholds were log transformed given their non-normal distributions in the reference data.

#### Assessment of heat pain sensitivity

A series of three 7-s stimuli, at 45, 44, and 46 °C, with a 20-s inter-stimulus interval, were delivered to the upper volar aspect of the right hand using a TSA-II thermode. Thermode temperature increased from baseline (32 °C) at a rate of 2 °C/s. Following each stimulus, participants were asked to rate their pain on a 0–100 Numeric Rating Scale. Thermode was slightly repositioned between stimuli to avoid adaptation/sensitization.

#### Pain 50 determination

Pain 50 was determined as the temperature at which a participant reported pain intensity of 50. If a pain rating of 50 was not reached during the heat pain sensitivity test, the stimulus temperature was increased or decreased (up to a maximal temperature of 48.5 °C) until a NPS rating of 50 was reached.

#### Conditioning stimulus

Participants were asked to place their left hand in a 12 °C water bath for 10 s, and then rate their pain intensity. If pain intensity fell below 40, the water temperature was decreased at 0.5 °C intervals down to a minimum of 10 °C.

#### Conditioned pain modulation (CPM)

Test-stimuli were applied as following: The individual pain-50 temperature was applied to the volar aspect of the right forearm for a duration of 30 s, and participants were asked to rate the pain intensity at 0, 10, 20, and 30 s.

Then, following a 10-min break, a conditioning stimulus was given as follows: participants were asked to immerse their left hand in a cold-water bath for 40 s and rate their pain intensity after 10 s. Immediately after rating the conditioning stimulus, the test stimulus was applied to the right volar area, as previously described, for 30 s while the left hand was still immersed in cold water. Participants were asked to rate the pain intensity elicited by the test stimulus at 0, 10, 20, and 30 s, and then to re-rate the pain elicited by the conditioning stimulus. CPM magnitude was calculated by subtracting the test-stimulus pain ratings at 30 s from the test + conditioning pain ratings at 30 s.

#### FMT procedure

Participants received a native microbiota depletion protocol^77^ that included a 3-day oral antibiotic regimen of vancomycin 500 mg (Mylan, France) and neomycin 1000 mg (Ma’ayan Haim, Israel), 24h of clear liquid diet and bowel irrigation using polyethylene glycol 255 g (PEG, Taro, Israel). Each participant received 30 healthy donor FMT capsules, the equivalent of 15 g. The induction dose of 30 capsules was followed by 4 maintenance doses of 15 capsules each, taken 2 weeks apart, for a total duration of 8 weeks.

#### Clinical follow-up

During the treatment period, participants were asked to fill in a daily symptom-diary, monitoring their level of pain, fatigue, and quality of sleep. They were contacted weekly and interviewed regarding their symptoms, change in regularly taken medications, and any side effects.

On each clinic visit, patients filled in the study questionnaires (as listed above in *baseline evaluation*). Venous blood was analyzed for determining HbA1C, lipid profile, liver enzymes, and renal function, at baseline and on weeks 1 and 4 post treatment.

## Acknowledgements

This study was supported by the Weston Family Foundation and The Louise and Alan Edwards Foundation to Y.S., A.M., and A.K.; W.C was supported by a postdoctoral fellowship from The Louise and Alan Edwards Foundation and the Fonds de recherche du Québec – Santé (FRQS).

## Author contributions

W.C., Y.S., A.M., and A.K. conceived the project, designed experiments, and supervised the research. W.C. performed behavioral and molecular studies in mice with the help of C.W., N.B., S.T., K.C.L., M.H., C.C., and N.W. J.Z., J.S.M., I.L.K., N.J.B.B., T.S., B.S., P.S., M.P.K., and E.G. assisted with study design and interpretation of results. E.G and N.J.B.B performed microbiome analyses. F.W., and Y.D.K. performed in vivo calcium imaging. I.K. and S.W. assisted with the gut analyses. M.H., R.H., I.K., T.H., R.L., H.B.Y., M.P. and A.M. performed clinical trial in humans and collected samples for mouse FMT. W.C., Y.S., A.M., and A.K. wrote the manuscript. All authors reviewed the manuscript and discussed the work.

## Data availability

Sequencing data generated in this study have been deposited in the Gene Expression Omnibus. 16S rRNA, WGS, and scRNAseq data were deposited under the BioProject accession number PRJNA1029994.

## Additional information

Supplementary Information is available for this paper. Correspondence and requests for materials should be addressed to Amir Minerbi and Arkady Khoutorsky.

